# Highly Genotype- and Tissue-specific Single-Parent Expression Drives Dynamic Gene Expression Complementation in Maize Hybrids

**DOI:** 10.1101/668681

**Authors:** Zhi Li, Peng Zhou, Rafael Della Coletta, Tifu Zhang, Alex B. Brohammer, Brieanne Vaillancourt, Anna Lipzen, Chris Daum, Kerrie Barry, Natalia de Leon, Cory D. Hirsch, C. Robin Buell, Shawn M. Kaeppler, Nathan M. Springer, Candice N. Hirsch

**Affiliations:** Department of Agronomy and Plant Genetics, University of Minnesota, Saint Paul, MN 55108; Department of Plant and Microbial Biology, University of Minnesota, Saint Paul, MN 55108; Provincial Key Laboratory of Agrobiology, Jiangsu Academy of Agricultural Sciences, Nanjing, China; Department of Plant Biology, Michigan State University, East Lansing, MI 48824; US Department of Energy Joint Genome Institute, Walnut Creek, CA 94598; Department of Agronomy, University of Wisconsin, Madison, WI 53706; Department of Plant Pathology, University of Minnesota, Saint Paul, MN 55108

## Abstract

Maize exhibits tremendous gene expression variation between different lines. Complementation of diverse gene expression patterns in hybrids could play an important role in the manifestation of heterosis. In this study, we used transcriptome data of five different tissues from 33 maize inbreds and 89 hybrids (430 samples in total) to survey the global gene expression landscape of F_1_-hybrids relative to their inbred parents. Analysis of this data set revealed that single parent expression (SPE), which is defined as gene expression in only one of the two parents, while commonly observed, is highly genotype- and tissue-specific. Genes that have SPE in at least one pair of inbreds also tend to be tissue-specific. Genes with SPE caused by genomic presence/absence variation (PAV SPE) are much more frequently expressed in hybrids than genes that are present in the genome of both inbreds, but expressed in only a single-parent (non-PAV SPE) (74.7% vs. 59.7%). For non-PAV SPE genes, allele specific expression was used to investigate whether parental alleles not expressed in the inbred line (“silent allele”) can be actively transcribed in the hybrid. We found that expression of the silent allele in the hybrid is relatively rare (∼6.3% of non-PAV SPE genes), but is observed in almost all hybrids and tissues. Non-PAV SPE genes with expression of the silent allele in the hybrid are more likely to exhibit above high-parent expression level in the hybrid than those that do not express the silent allele. Finally, both PAV SPE and non-PAV SPE genes are highly enriched for being classified as non-syntenic, but depleted for curated genes with experimentally determined functions. This study provides a more comprehensive understanding of the potential role of non-PAV SPE and PAV SPE genes in heterosis.

## Introduction

Modern maize production has benefited from heterosis, a widely observed phenomenon in which hybrids show superior performances relative to their inbred parents (Flint-Garcia et al., 2009; Birchler et al., 2010). However, the manifestation of heterosis has been shown to vary greatly across traits and environments for a given hybrid (Flint-Garcia et al., 2009; Li et al., 2018), making it unlikely to be explained by a single mechanism. Indeed, the molecular underpinnings for heterosis are still under debate (Schnable and Springer, 2013; Wallace et al., 2018). Over the past century, various models including dominance, overdominance, and epistasis have been proposed to explain heterosis (Shull, 1908; Bruce, 1910; Jones, 1917; East, 1936; Birchler et al., 2010). Nevertheless, it is still unclear how the combination of two sets of parental alleles leads to hybrid vigor.

The advent of next-generation sequencing has enabled large-scale exploration of genomic variation and has found extensive sequence variation in maize (Buckler et al., 2006; Gore et al., 2009; Chia et al., 2012; Romay et al., 2013; Bukowski et al., 2018). Studies using genome wide association study (GWAS) and quantitative trait locus (QTL) mapping have identified numerous loci associated with phenotypic variation. However, many studies support that sequence diversity is necessary but not sufficient to generate heterotic phenotypes (Kaeppler, 2012). In addition to sequence variation, maize genomes exhibit high levels of structural variation, such as copy number variation (CNV) and presence/absence variation (PAV), which is also a potential contributor to phenotypic variation (Springer et al., 2009; Swanson-Wagner et al., 2010; Jiao et al., 2012; Maron et al., 2013; Hirsch et al., 2014; Hirsch et al., 2016; Brohammer et al., 2018). For example, the copy number of the *Mate1* gene is positively associated with seedling aluminum tolerance in maize roots (Maron et al., 2013). Together, this high level of genomic variation has led to the hypothesis that the two sets of parental alleles might act synergistically in the hybrid to generate heterosis (Fu and Dooner, 2002; Lai et al., 2010; Sun et al., 2018).

Differential gene expression has also been suggested to play a vital role in shaping plant phenotypes (Harrison et al., 2012; Wallace et al., 2014). Substantial variation has been observed in maize transcriptomes among diverse inbred lines (Paschold et al., 2012; Hansey et al., 2012; Fu et al., 2013; Hirsch et al., 2014; Kremling et al., 2018). Efforts have been made to investigate heterosis at the level of transcriptome differentiation between inbreds and hybrids (Swanson-Wagner et al., 2006; Springer and Stupar, 2007a). Several hypotheses have been proposed to explain heterosis at the level of the transcriptome, such as a possible role of additive (Stupar and Springer, 2006; Swanson-Wagner et al., 2006; Guo et al., 2006; Meyer et al., 2007; Springer and Stupar, 2007a; Stupar et al., 2008; Thiemann et al., 2010; Thiemann et al., 2014) or nonadditive gene expression (Auger et al., 2005; Użarowska et al., 2007; Pea et al., 2008; Hoecker et al., 2008; Swanson-Wagner et al., 2009; Jahnke et al., 2010; Paschold et al., 2010; Baldauf et al., 2016), as well as the interaction of the two parental alleles in hybrids (Springer and Stupar, 2007b).

More recently, the role of single parent expression (SPE) genes in heterosis has been investigated (Paschold et al., 2012; Paschold et al., 2014; Baldauf et al., 2016; Marcon et al., 2017; Baldauf et al., 2018). SPE refers to a situation in which expression can only be detected in one of the two parental lines of a hybrid. These studies have shown that more genes are expressed in hybrids relative to either parent via SPE gene expression complementation which may contribute to heterosis. SPE patterns could arise solely due to expression differences (on/off) between the two shared parental alleles (non-PAV SPE) or as a result of the physical absence of the gene in one of the parental inbred lines (PAV SPE). The distinction between PAV SPE and non-PAV SPE has yet to be investigated and it remains unclear whether these categories of SPE lead to differences in hybrid expression complementation across diverse tissues and genetic backgrounds.

In this study, transcriptome data from 89 hybrids derived from 33 maize inbreds in five diverse tissues was used to document the frequency and impact of single parent expression on hybrid expression patterns across diverse genotypes and tissues. We partition single parent expression into instances where the gene is present in both parental lines but only expressed in one parent (non-PAV SPE) and when the gene is present in only one of the parental lines (PAV SPE). Results of this study shed light on the potential role of non-PAV SPE and PAV SPE genes in the manifestation of heterosis.

## Results

### Transcriptome Profiling of a Diverse Panel of Maize Inbred Lines and Their F_1_ Hybrids in Five Different Tissues

We selected 33 diverse maize inbred lines with representation across the major heterotic groups in U.S. corn breeding programs to assess transcriptome differences in inbred parents vs F_1_ hybrids (Figure 1A). Five of the inbreds (B73, Mo17, PH207, Oh43 and PHG29) were used as male parent crossing with other inbreds to generate F_1_ hybrids. A total of 89 different hybrids were generated, including 30 inbreds crossed by B73, 30 inbreds crossed by Mo17, 27 inbreds crossed by PH207, one inbred crossed by Oh43, and one inbred crossed by PHG29. RNA-sequencing (RNA-Seq) was performed on all of the inbred parents and F_1_ hybrids in five distinct tissues (seedling root and shoot at the V1 developmental stage, leaf and internode at the V7/V8 development stages, and endosperm at 15 days after pollination (DAP); Supplemental Figure 1). A total of 430 RNA-Seq samples including 160 inbred and 270 hybrid samples across the five tissues were collected. Read numbers ranged from 20.9 to 51.7 (33.9 on average) million per sample (Supplemental Table 1). Tissues were clearly distinguished from one another based on transcript abundance by principal component analysis (PCA), and inbred and hybrid samples separated from each other for leaf, root, internode and seedling samples, but not endosperm (Figure 1B).

**Figure 1.**
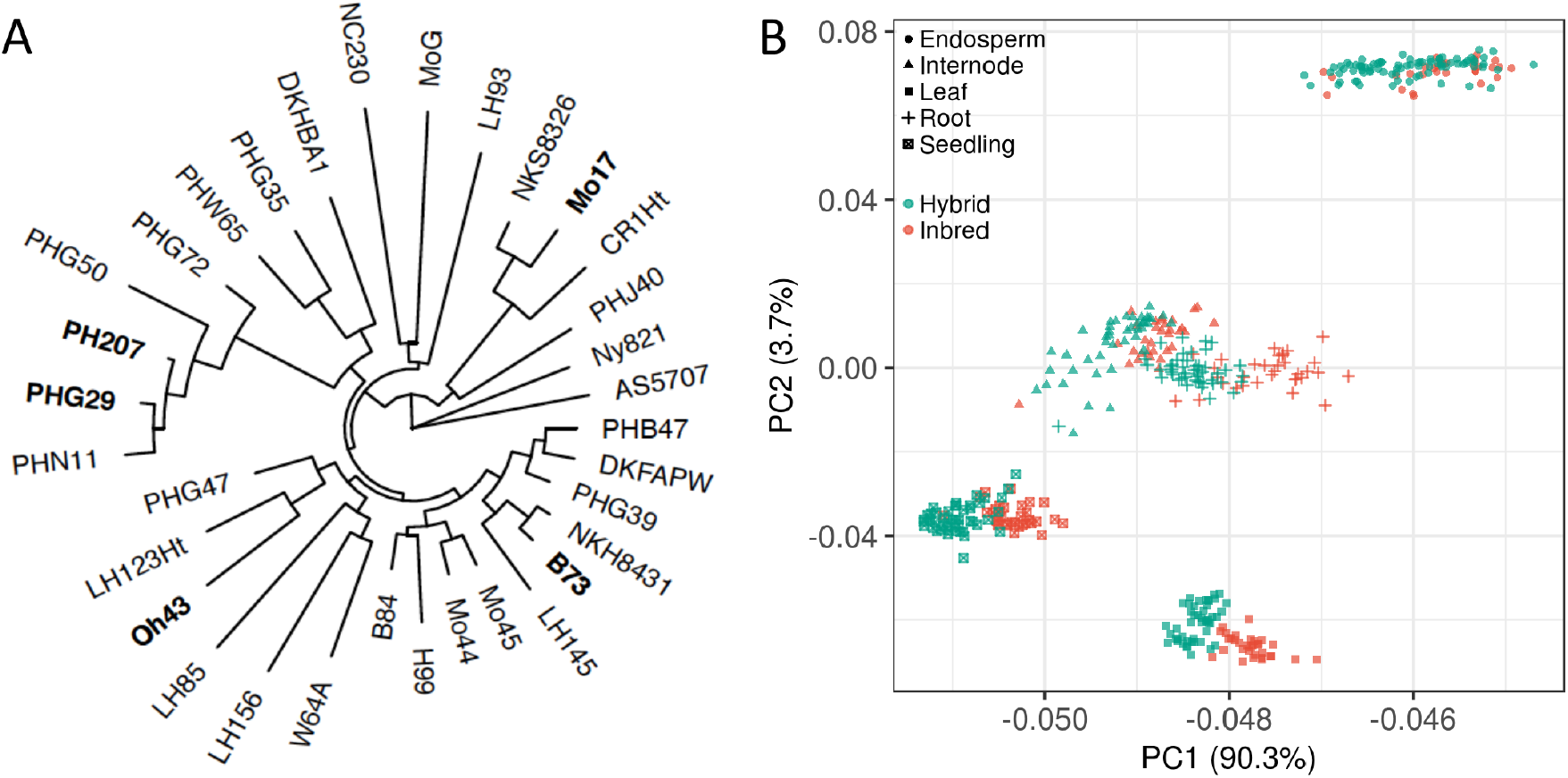
Genetic distance and transcriptomic relationship of maize inbred lines used in this study. (A) Maximum likelihood tree based on 532,329 SNPs of inbred parents in this study. The five male inbred parents of the hybrids are highlighted in bold (B73, PH207, Mo17, Oh43, PHG29). Due to the small number of genotypes included in the tree, groupings may not reflect precise genetic relationships. (B) Principal component analysis (PCA) of gene expression for 160 inbred and 270 hybrid samples across five tissues. Genes expressed in at least one sample were used for PCA. Shapes indicate different tissues and colors indicate hybrids versus inbreds.

Of the 39,324 predicted protein coding genes in the B73v4 reference genome, 31,502 genes (80.1%) were expressed (counts per million (CPM) >= 1) in at least one sample, with 16,560 to 22,215 genes expressed in each sample. We found 7,822 genes (19.9%) were not expressed (CPM=0) in any sample or expressed at very low levels (CPM< 1), while 10,137 genes (25.8%) were constitutively expressed in all 430 samples. Substantial variation in the number of expressed genes was observed among genotypes and tissues, with endosperm samples having the lowest number of genes expressed (17,340 on average) and root samples having the most genes expressed (21,048 on average).

We next characterized the number of expressed genes shared among different sets of samples and tissues. Similar trends in the frequency of expression across genotypes was observed for inbred or hybrid samples from the same tissue (Figure 2A and B). On average across the five tissues, 13,992 (35.6%, 12,015-15,643) genes were not expressed in any inbred or hybrid samples of a certain tissue (designated “Silent”); 3,863 (9.8%, 3,103-4,445) genes were expressed in less than 20% of the genotypes (designated “Genotype specific”), 4,048 (10.3%, 3,283-4,764) genes were expressed in 20%-80% of the genotypes (designated “Intermediate frequency”), and 17,421 (44.3%, 15,018-19,299) genes were expressed in more than 80% of the genotypes (designated “Constitutive”).

**Figure 2.**
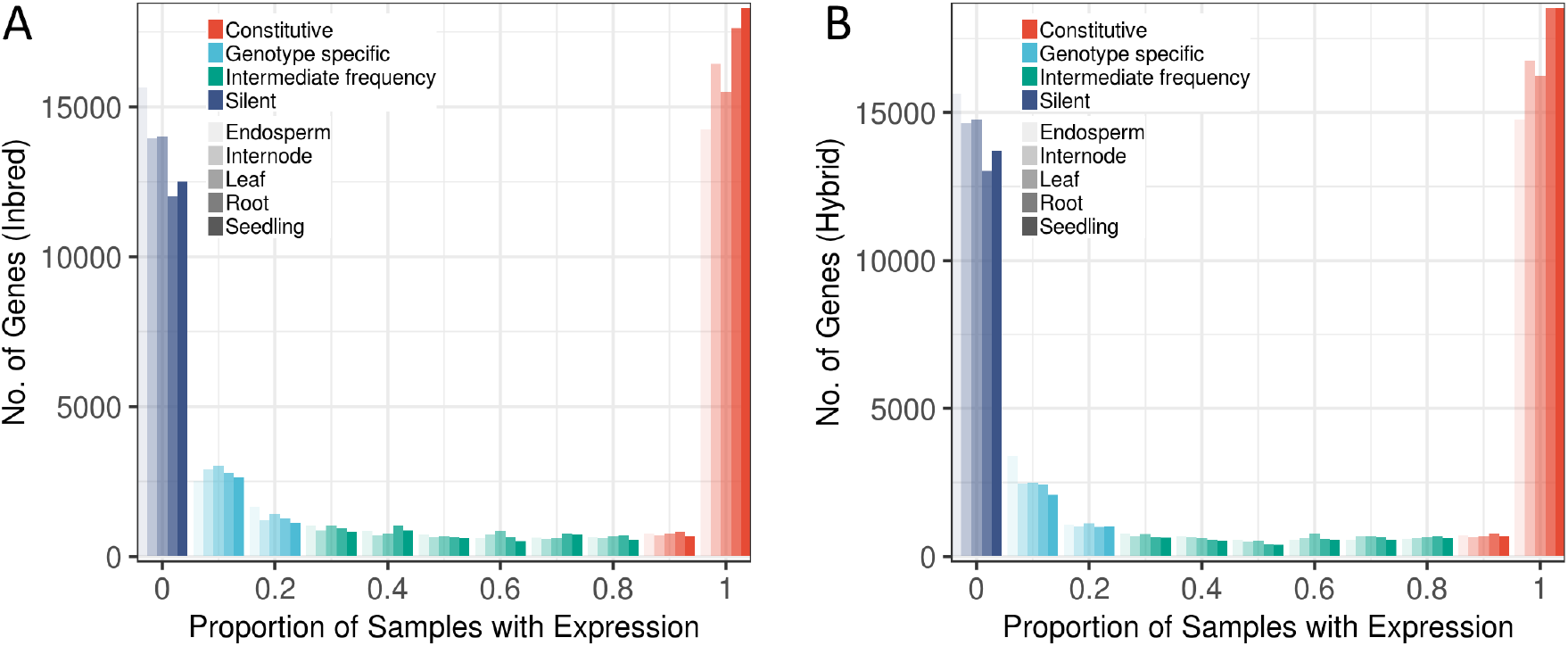
Summary of the gene expression landscape in diverse inbreds and hybrids across five tissues. (A, B) The number of genes detected across all available samples for inbreds (A) and hybrids (B) in each tissue (Endosperm, 30/88; Internode, 32/43; Leaf, 32/48; Root, 33/48; Seedling, 33/48, where the number before the slash is the inbred number and the number after the slash is the hybrid number). The transparency of bars represents different tissues, and colors represent the proportion out of all samples in each tissue that each gene is expressed (not expressed in any sample of that tissue (“Silent”), expressed in less than 20% of samples (“Genotype specific”), expressed in 20%-80% of samples (“Intermediate frequency”) and expressed in more than 80% of samples (“Constitutive”)).

### Hybrids do not Always Express More Genes Than Their Inbred Parents

To test the hypothesis whether hybrid genotypes express more genes than their inbred parents, 266 parent-hybrid triplets were constructed (Supplemental Figure 1E), and the number of expressed genes in the inbred parents and the hybrids were compared. Although generally hybrids expressed significantly more genes than inbreds in all tissues except seedling (139 to 427 more genes expressed in hybrids across tissues, t test, *P* < 0.05) (Supplemental Figure 2), there are exceptions (Figure 3, Supplemental Figure 3). Indeed, in 20% (57 out of 266) of the triplets, the hybrid expressed fewer genes than the average number of genes expressed in the two inbred parents, especially in seedling samples where over half of the triplets show this trend. To test if this was due to the threshold (i.e., CPM > 1) used to determine if a gene is expressed, four additional thresholds were tested (CPM of 0.5, 1.5, 2.0, and 3.0) and the results were consistent (Supplemental Figure 4).

**Figure 3.**
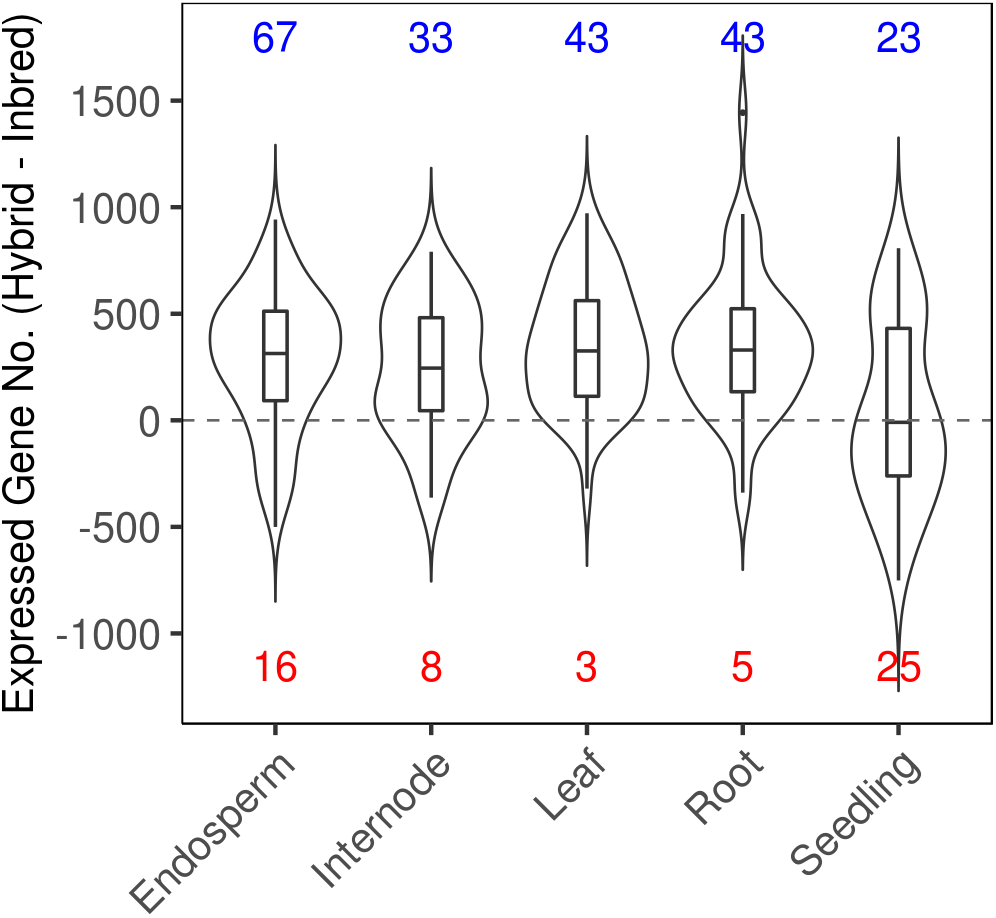
Difference of the number of expressed genes for each parent-hybrid triplet calculated as Hybrid - 0.5*(Male-parent + Female-parent) for each tissue. Red numbers indicate the number of parent-hybrid triplets in which there are more genes expressed in the parent mean, and blue numbers indicate the number of parent-hybrid triplets in which there are more genes expressed in the hybrid.

### Most PAV Genes are Not Expressed

Genes with presence/absence variation were previously called for each parental inbred by mapping resequencing reads to both the B73 and PH207 genomes (Brohammer et al., 2018). In total, 4,277 B73 genes were identified as putatively absent in at least one inbred, while 5,931 PH207 genes were identified as putatively absent in at least one inbred (Supplemental Figure 5A). Presence/absence variants relative to the PH207 genome assembly were filtered to retain only those that were syntenic to genes within the B73 reference genome assembly. After this filtering and removing genes with contradictory expression support (i.e. called putatively absent from the resequencing data and expressed in the inbred parent), a total of 4,060 B73 based PAV genes (PAV_B; 422-1,059 across 25 genotypes) and 1,493 PH207 based PAV genes (PAV_P; 67-265 across 25 genotypes) were retained (Supplemental Figure 5B, Supplemental Table 2, 3, 4).

We were interested to evaluate the general expression level of the PAV genes. To do this, we first classified the PAV genes based on their expression patterns into those that were expressed in at least one tissue in B73 or PH207 (ie., PAV SPE) and those that were not expressed in any tissue (PAV_Off; Figure 4). Only 14.6% (592) of PAV_B genes and 13.1% (196) of PAV_P genes were expressed in the B73 or PH207 inbred in at least one tissue (denoted as PAV_B SPE and PAV_P SPE respectively; Figure 5A; Supplemental Figure 6), a much lower rate compared with the full gene set (∼48.9%, 19,214/39,324). The number of PAV_B SPE and PAV_P SPE genes also varied significantly across genotypes and tissues (Figure 5B, Supplemental Figure 7A). Most PAV SPE genes were limited to specific genotypes and/or tissues. Only 5.7% of PAV SPE genes were shared in over half of all genotypes, while 67% were detected in less than 20% of all genotypes (Figure 5C, Supplemental Figure 7B). Additionally, 47.5% of PAV SPE genes were specific to one or two tissues and 25.8% were expressed in all five tissues (Figure 5C). In contrast, for non-PAV genes in B73 or PH207, ∼56.4% were shared in all five tissues, and only ∼20.3% were specific to one or two tissues (Supplemental Figure 8). Furthermore, we found that genotype-specific PAV_B SPE genes (blue bars in Figure 5C) also tended to be tissue-specific (blue pie-chart in Figure 5C).

**Figure 4.**
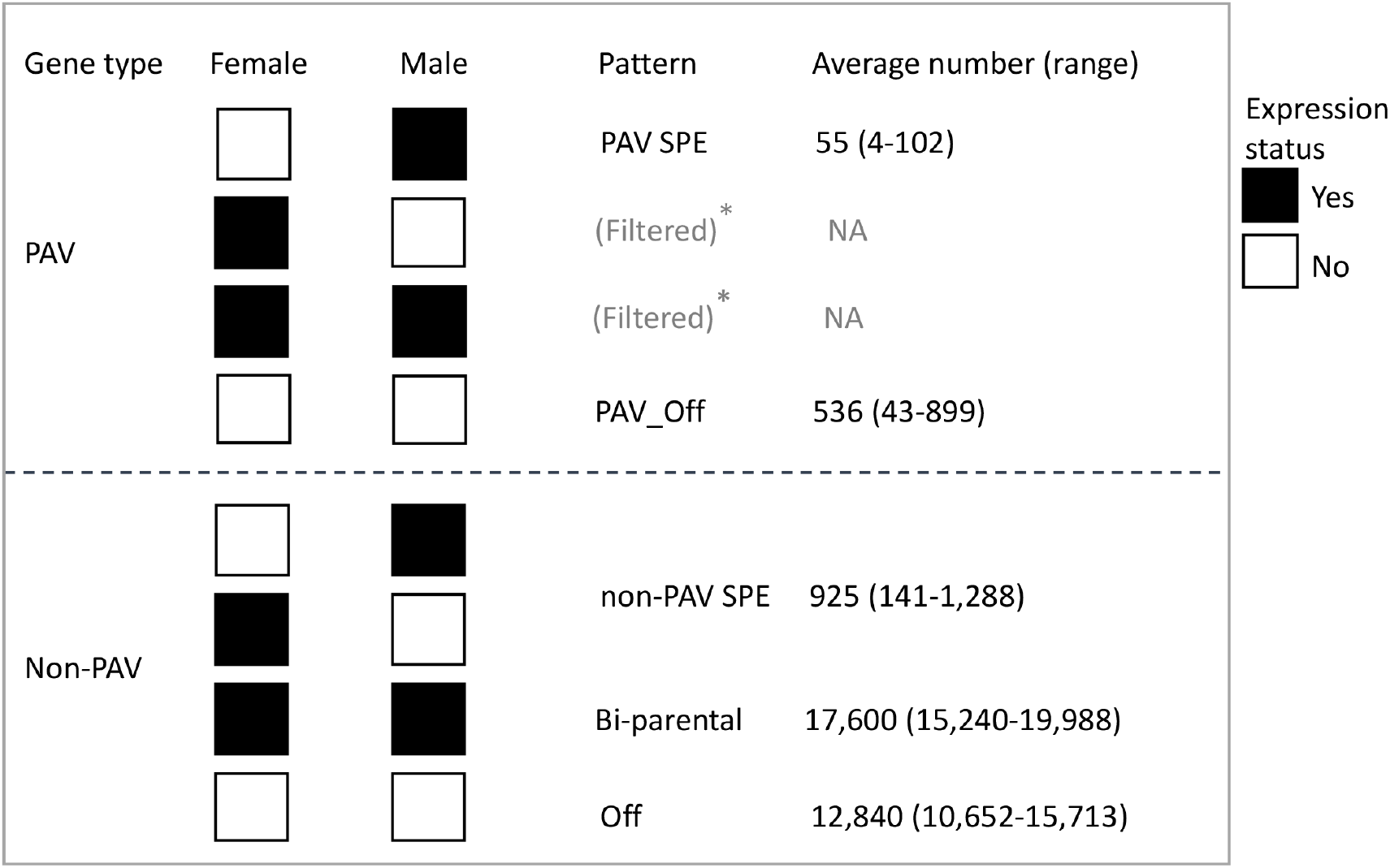
Criteria used to assign each gene into different expression patterns. The (*) indicates patterns that were filtered from downstream analyses. Range is the range of gene numbers for each pattern observed across the tissues and genotypes.

**Figure 5.**
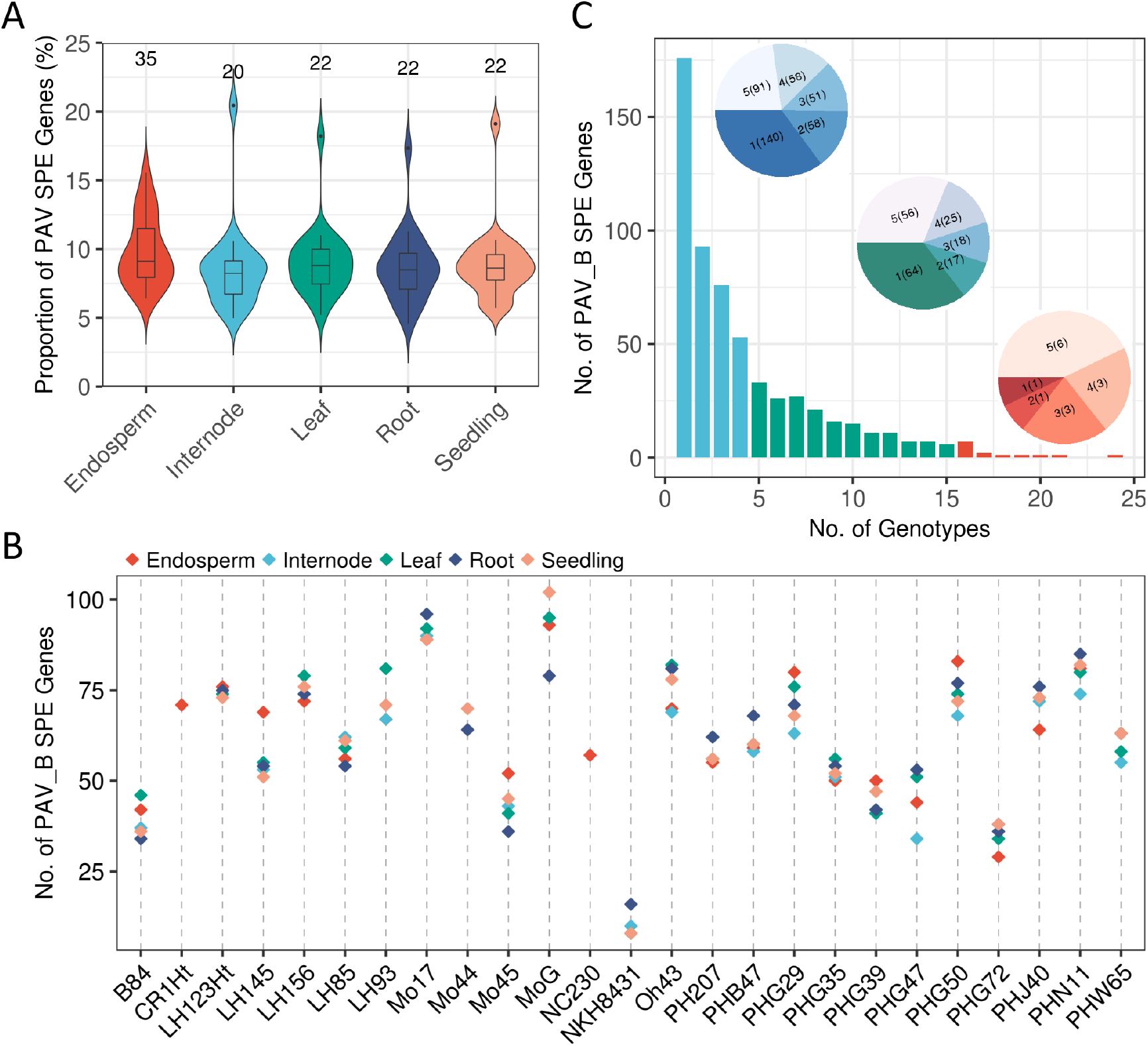
Summary of PAV SPE genes. (A) The proportion of PAV genes expressed in the B73 (PAV_B SPE) or PH207 (PAV_P SPE) genome out of all PAV genes identified across the different genotypes. (B) The number of PAV_B SPE genes in each female parent and tissue. (C) The number of PAV_B SPE genes detected in 1-25 female parents and 1-5 tissues. Light blue, green, and red color bars represent the number of PAV_B SPE genes shared across <20%, 20%-80%, and >80% of the total inbreds, respectively. The embedded pie chart with similar color scale of the corresponding group of bars shows the PAV_B SPE gene numbers expressed in 1 to 5 tissues of that bar group. Each part of the embedded pie chart represents the number of PAV_B SPE genes (number in parentheses) shared by the number of tissues (1-5, outside parentheses).

### Single Parent Expression is Common and Highly Genotype-specific and Tissue-specific

Aside from PAV SPE, SPE can also be caused by expression differences (on/off) between two shared parental alleles (non-PAV SPE). In this study we required a gene in one parent to have a CPM >=1 and the same gene in the other parent to have a CPM < 0.1 to be classified as non-PAV SPE genes. In total, 12,808 non-PAV SPE genes were identified across all 266 triplets, ranging from 146 to 1,309 per triplet. There was substantial variation in the number of non-PAV SPE genes among genotypes and tissues (Figure 6A). We found that if the genetic distance between two parents is relatively low, the number of non-PAV SPE genes identified between them also tended to be low. For example, four of the inbreds PHG29, PHG50, PHG72 and PHN11 are closely related with the common paternal inbred PH207. The number of non-PAV SPE genes identified between these inbreds and PH207 were among the lowest of all tested triplets. Similar observations were observed with parental pairs such as NKH8431 vs. B73, LH145 vs. B73, NKS8326 vs. Mo17 and CR1Ht vs. Mo17 (Figure 6A). Similar to PAV SPE genes, non-PAV SPE genes were also expressed in a highly genotype- and tissue-specific manner compared with the full gene set, where nearly half of the genes were expressed in over 80% of the genotypes (Figure 2), and ∼56.4% genes were expressed in all five tissues (Supplemental Figure 9). We observed only ∼1.6% (200) non-PAV SPE genes with expression in over 50% of the triplets and no non-PAV SPE gene was expressed in over 80% of the triplets (Figure 6B). In addition, ∼66% of the non-PAV SPE genes were specific to only one or two tissues, and only ∼16.3% showed non-PAV SPE in all five tissues (in at least one triplet). For non-PAV SPE genes specific to less than 20% of the genotypes, the majority of them were also specific to a single tissue (Figure 6B).

**Figure 6.**
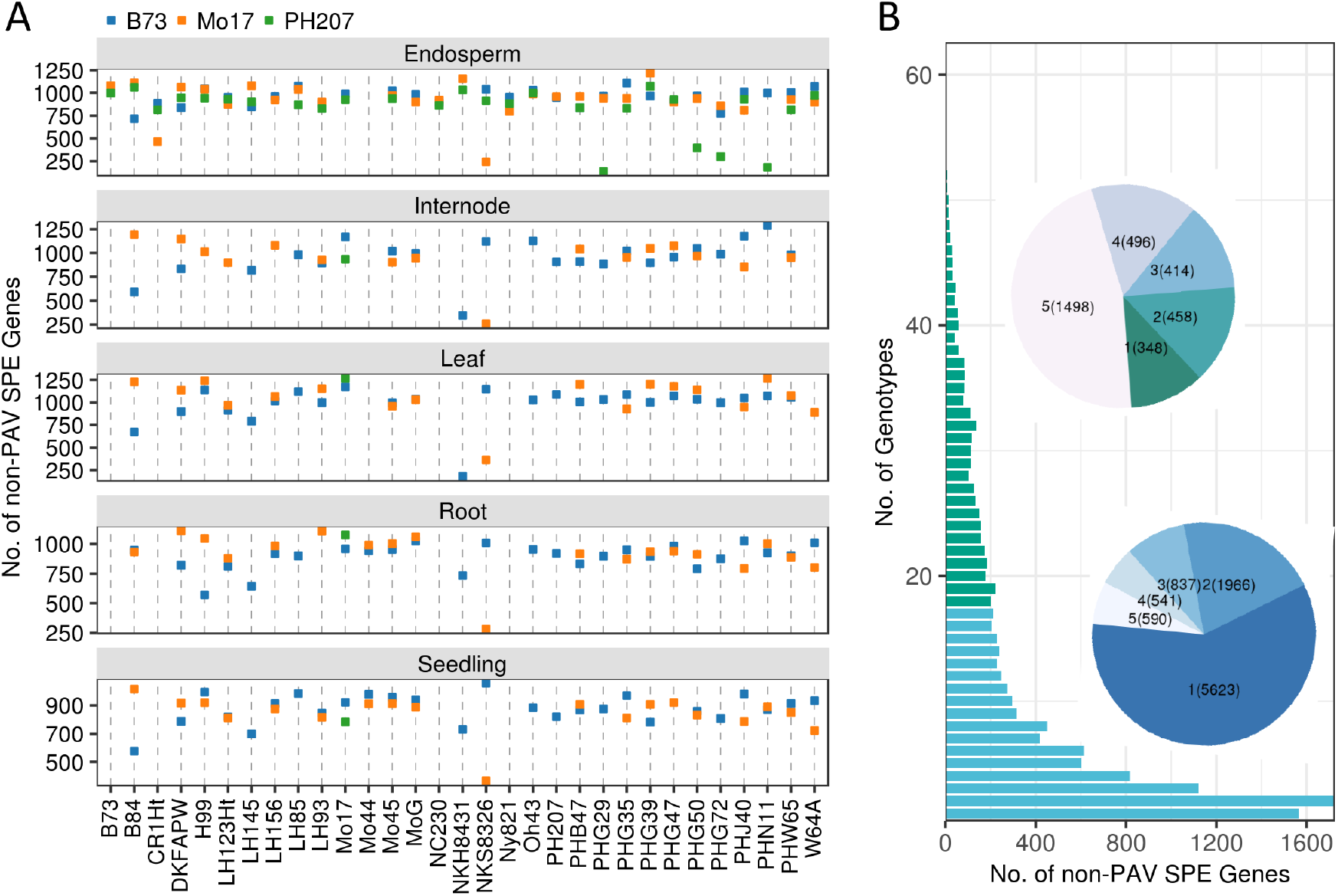
The distribution of non-PAV SPE genes across different genotypes and tissues. (A) The number of non-PAV SPE genes in each pair of inbred parents and tissue. Different colors indicate the three common male inbred parents. (B) The number of non-PAV SPE genes detected in 1-30 inbred parent pairs and 1-5 tissues (embedded pie chart). Light blue, green, and red color of bars represent the number of non-PAV SPE genes shared across <20%, 20%-80%, and >80% of total inbred parent pairs, respectively. The embedded pie chart with similar color scale of the corresponding group of bars shows the expressed non-PAV SPE gene numbers in 1 to 5 tissues of that bar group. Each part of the embedded pie chart represents the number of non-PAV SPE genes (number in parentheses) shared by the number of tissues (1-5, outside parentheses).

### PAV SPE and non-PAV SPE Genes Show Distinct Patterns of Expression Complementation in Hybrids

We were interested to know the extent of expression complementation that is observed in hybrids for PAV SPE and non-PAV SPE genes as this could be a mechanism that contributes to the superior performance of hybrids relative to their inbred parents. Genes silent in both parents rarely showed expression in the hybrid (average of ∼4.3% across triplets; Figure 7A, Supplemental Table 5 “PAV_Off” and “Off” categories). Likewise, the vast majority of genes expressed in both parents (i.e., “Bi-parental”) were expressed in the hybrid (98.0%). For non-PAV SPE genes, 59.7% were expressed in the hybrid. Interestingly, a significantly higher proportion of PAV SPE genes (i.e., genomic PAV genes with parental expression support) were expressed in the hybrid compared with non-PAV SPE genes (74.7%, Wilcoxon rank-sum test, *P* < 2.2e-16; Figure 7A). However, the fraction of non-PAV SPE and PAV SPE genes expressed in the hybrids varied substantially across genotypes (28.6%-79.7% for non-PAV SPE genes, 43.5%- 94.9% for PAV SPE genes) and across tissues (Supplemental Table 5). The expression level of non-PAV SPE and PAV SPE genes was similar to genes classified as bi-parental, except for a lack of highly expressed genes (log2(CPM) >10; Supplemental Figure 10A-C). Thus, the observed differences between non-PAV SPE and PAV SPE genes did not seem to be due to an excess of lowly expressed genes. Taken together, the significant difference observed between non-PAV SPE and PAV SPE genes for the proportion of genes expressed in hybrids may indicate distinct roles in hybrid expression complementation. It should be noted however, due to the lack of reference genomes for every genotype in this study, we cannot identify reciprocal PAV genes between each pair of the two inbred parents. We therefore likely underestimated the amount of PAV and PAV SPE genes in these triplet combinations. Thus, the impact of PAV SPE genes on hybrid expression complementation may be even more significant if these non-B73 PAV genes were included.

**Figure 7.**
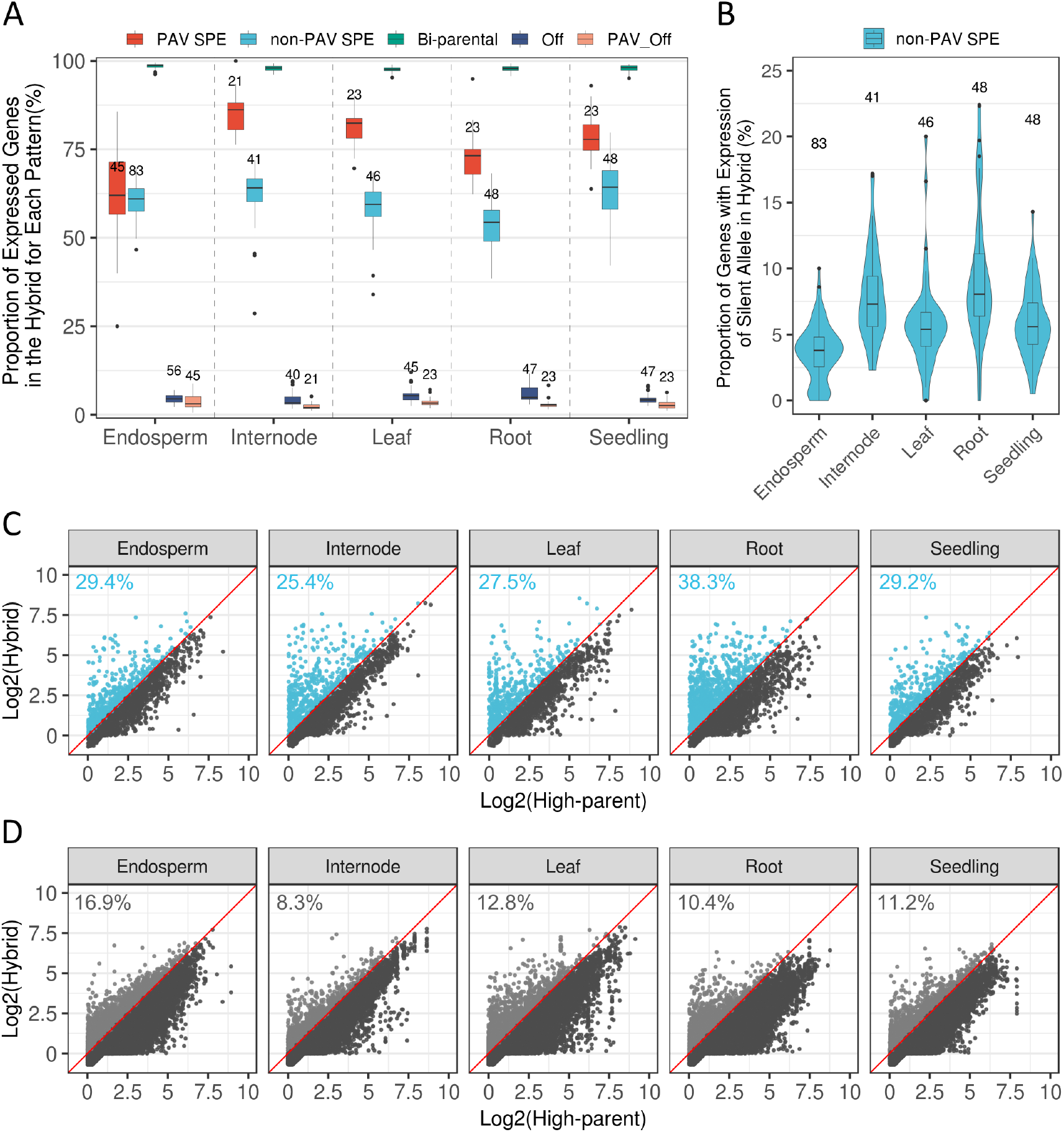
Silent allele expression in the hybrid of non-PAV SPE genes. (A) The proportion of genes expressed in the hybrid for each pattern for 266 triplets. The number of parent-hybrid triplets of each tissue for each pattern is shown above each box. (B) The proportion of silent allele expression in hybrids for non-PAV SPE genes. The number of parent-hybrid triplets of each tissue is shown above each box. (C, D) Expression level of non-PAV SPE genes with (C) and without (D) silent allele expression in the hybrid. The log2-transformed expression level (CPM) of each gene in the high-parent inbred and the hybrid was plotted. Dots that fall above the red line indicate genes that have above high-parent (AHP) expression level in the hybrid. Numbers in each panel indicate the proportion of genes with AHP expression level in the hybrid.

### Expression of Parentally Silent Alleles Occurs in Most Hybrids, but at a Low Frequency within Each Hybrid

We observed expression of many non-PAV SPE genes in the hybrids in this study. To further investigate the nature of hybrid expression of the SPE genes, we assessed the relative expression of the two parental alleles in the hybrid using allele-specific expression (ASE) data. On average, over 4.7 million SNPs were identified for each of the 30 inbred parents relative to the B73 reference genome using 20-40x depth resequencing data, and these SNPs were used to quantify expression levels of each allele in the hybrids. On average across the triplets, 562 non-PAV SPE genes had sufficient coverage in at least two SNP positions in the hybrid and could be assayed for ASE, accounting for 59.9% of the total non-PAV SPE genes for each hybrid (Supplemental Table 5). Evidence of expression (allele-specific reads > 10) was detected for the parentally silent allele for 2,912 non-redundant non-PAV SPE genes, and the result was robust when applying different cutoffs (Supplemental Figure 11). We found expression of the silent allele from at least one non-PAV SPE gene occurred in almost all hybrids (97.7%, 260/266). However, the proportion of genes in which the silent allele was expressed in the hybrid was quite low (average of 35 genes or 6.3% of all non-PAV SPE genes), although there was substantial variation across genotypes and tissues (Figure 7B). In addition, when comparing the expression level of SPE genes in the hybrid with the high-parent value, we found that non-PAV SPE genes with silent allele expression in the hybrid were more likely to show above high-parent expression levels compared to those without expression of the silent allele in the hybrid (30.4% vs. 12.7%; Figure 7C, D, Supplemental Figure 12).

### PAV SPE and non-PAV SPE Genes Are Highly Enriched in Non-syntenic Genes and Depleted in Genes with Experimentally Determined Function and/or Literature Support

Single parent expression genes (including both PAV SPE and non-PAV SPE) are by definition non-essential genes as they are required to be present or expressed in the tissues of only some inbred lines. Furthermore, the number of PAV SPE and non-PAV SPE genes is highly variable across tissues and genotypes and may contribute to very specific functions within certain tissues or environmental conditions. To examine the broad functionality of these genes, we utilized two sources of gene classifications. The first classification we tested was genes that are orthologous to sorghum (syntenic), versus those that are non-syntenic. Of the 39,324 high confidence protein coding genes in the B73v4 reference genome, ∼39.2% (15,402/39,324) did not have an identifiable syntenic ortholog in the sorghum ancestor genome (i.e., non-syntenic; (Brohammer et al., 2018). Both PAV SPE and non-PAV SPE genes were highly enriched for being non-syntenic, with PAV SPE genes having an even higher proportion of genes categorized as non-syntenic on average (83.1%) than non-PAV SPE genes (78.1%). In contrast, genes showing a “Bi-parental” pattern of expression were consistently depleted (average 13.1%) in genes categorized as non-syntenic across genotypes and tissues (Figure 8A; Supplemental Table 5).

**Figure 8.**
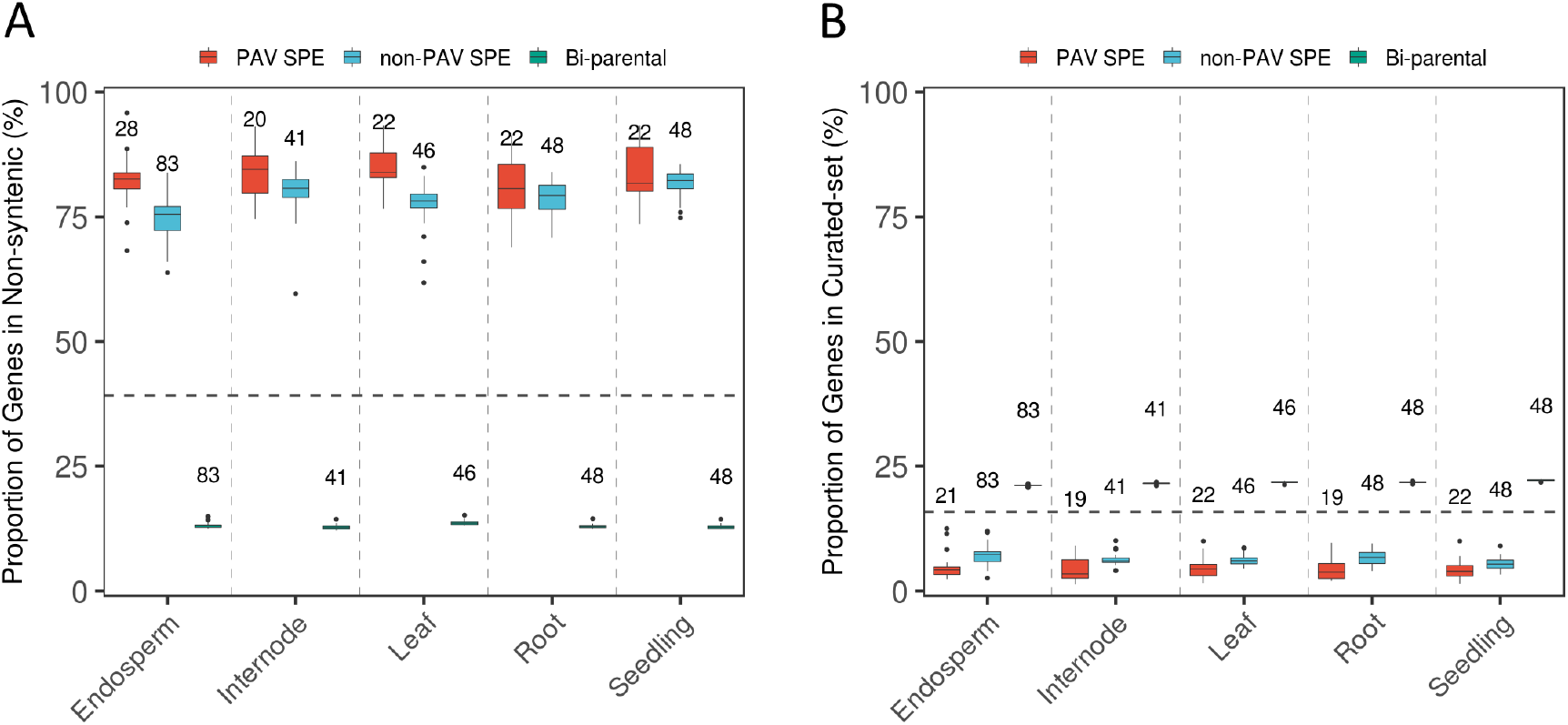
Proportion of genes in the non-syntenic and curated gene sets for each expression pattern. (A) Proportion of genes from each expression pattern that are in the non-syntenic gene list. Dashed black line represents the genome-wide proportion of non-syntenic genes (39.2%, 15,402/39,324). (B) Proportion of genes in each expression pattern that are in the curated gene set. Dashed black line represents the genome-wide proportion of curated genes (15.9%, 6,239/39,324). The number of parent-hybrid triplets of each tissue for each pattern is shown above each box. To avoid the inflation of proportions caused by a small number of genes in each pattern, only triplets with gene numbers greater than 20 in each pattern were included in this analysis.

We also tested enrichment of a set of 6,239 genes with experimentally determined function and/or literature support that had been previously curated (Andorf et al., 2016), and includes 15.9% (6,239/39,324) of all protein-coding genes in the B73v4 genome. We found the proportion of curated genes in each expression pattern was opposite to the proportions observed for non-syntenic genes. Compared to the genome-wide fraction of curated genes (15.9%), PAV SPE and non-PAV SPE genes were depleted (6.4% and 4.5% respectively) in the curated gene set, while genes classified as Bi-parental were enriched for curated genes across samples (average 21.6%; Figure 8B; Supplemental Table 5).

## Discussion

Transcriptome divergence between maize inbred parents and their F_1_-hybrid progeny has been previously investigated (Guo et al., 2004; Swanson-Wagner et al., 2006; Springer and Stupar, 2007b; Stupar et al., 2008; Paschold et al., 2012; Paschold et al., 2014; Baldauf et al., 2016; Marcon et al., 2017; Waters et al., 2017; Baldauf et al., 2018). While these studies provided insights into transcriptional variation between inbreds and hybrids and its potential role in heterosis, they were performed in one or a handful of parent-hybrid triplets or only a single tissue. In this study we constructed 266 parent-hybrid triplets across diverse heterotic backgrounds and tissues. This study provides a more comprehensive view of transcriptome divergence between inbred parents and hybrids and how this variation could potentially relate to hybrid vigor than has been previously documented.

We examined the number of expressed genes in hybrids and their inbred parents and found that the differences were highly tissue-dependent and genotype-dependent. In a previous study using maize root tissue of different developmental stages it was shown that hybrids consistently expressed more genes than inbred parents in all the investigated triplets (Paschold et al., 2012; Paschold et al., 2014; Baldauf et al., 2018). In this study, this same trend was observed for the vast majority of the root triplets (90%, 43 out of 48) (Figure 3). Furthermore, triplets (or reciprocal hybrids, Mo17 x B73, Oh43 x B73, H99 x B73, W64A x B73) examined by both studies exhibited very consistent results (Supplemental Figure 3). Another recent study using a maize developmental transcriptome (23 tissues) in B73, Mo17, as well as their F_1_ hybrid (Zhou et al., 2019) showed the same trend in most of the investigated tissues (83%, 19 out of 23), including all three root tissues. However, in our study, which included triplets with more genotype x tissue combinations, we found the opposite pattern (i.e., the average number of genes expressed in the parental inbreds was higher than in the hybrid) in over 20% of the 266 triplets (Figure 3). The difference in expressed gene number increased as the minimum CPM threshold increased (Supplemental Figure 4). We further assigned each gene into five expression patterns (Figure 4), and evaluated the hybrid expression profile for each category. We found 59.7% of non-PAV SPE and 74.7% of PAV SPE genes are expressed in hybrids and contribute to hybrid genotypes expressing more genes in general than their inbred parents.

We detected expression of silent alleles in almost all of the inbred-hybrid triplets (97.7%, 260/266). However, the proportion of silent alleles expressed in any individual hybrid was low (average 35 genes or 6.3% of the total non-PAV SPE genes on average in the triplet). These results imply that non-PAV SPE genes are predominantly under cis-regulation, although trans-regulation can occassionally occur. Interestingly, non-PAV SPE genes with silent allele expression in the hybrid are more likely to show above high-parent (AHP) expression levels in the hybrid compared with those without expression of the silent allele in the hybrid (30.4% vs. 12.7%), and this rate is similar to the AHP fraction for bi-parental expressed genes (Figure 7C, Supplemental Figure 10, 11). This result indicates that non-PAV SPE genes with silent allele expression in the hybrid are probably under different inheritance pattern (i.e., non-additive or dominant) from those without silent allele expression in the hybrid, which potentially contributes to heterosis.

Maize experienced its last whole genome duplication event ∼5-12 million years ago, right after the divergence of maize and sorghum (Swigoňová et al., 2004). Even though the subsequent diploidization process occurring in the tetraploid progenitor of maize caused dramatic genome rearrangements and fractionation (Woodhouse et al., 2010; Schnable, 2015), large blocks of syntenic regions can still be found between maize and sorghum (Schnable et al., 2011a). A recent study showed that about 60% of the total high confident protein-coding genes in the B73v4 genome have a syntenic ortholog in sorghum (Brohammer et al., 2018). Syntenic genes in the maize genome are thought to be more ancient, stable, and related to important biological functions. In contrast, non-syntenic genes are more likely the result of subfunctionalization and neofunctionalization and less related to visible phenotype (Schnable and Freeling, 2011; Schnable, 2015). For the 10,137 genes expressed in all 430 samples in this study, over 90% were found in maize-sorghum syntenic blocks and annotated with potential functions (curated gene set) (Supplemental Figure 13). These results indicate a vital role of constitutively expressed syntenic genes for the maintenance of basic cellular function. In contrast, we found PAV SPE and non-PAV SPE genes are highly enriched in the non-syntenic gene space and depleted for genes with possible functions from the curated gene set, consistent with previous findings (Swanson-Wagner et al., 2010; Paschold et al., 2014; Marcon et al., 2017; Baldauf et al., 2018).

Non-syntenic genes have been proposed to contribute to the adaptation of plants to fluctuating environments. For example, disease resistance genes were found to be enriched in the non-syntenic portion of the *Arabidopsis thaliana* (Freeling et al., 2008), *Oryza sativa* (Xu et al., 2011), and *Aegilops tauschii (Dong et al., 2016)* genomes. Non-syntenic genes may also play an important role in the ability of hybrids to deal with abiotic stress conditions (Marcon et al., 2017). While syntenic and non-syntenic genes seem to be specialized in different aspects of plant life, the interaction of syntenic and non-syntenic genes in different developmental stages can have an important impact on phenotype. Furthermore, the enrichment of non-syntenic genes in non-PAV SPE and PAV SPE gene sets, along with the enrichment for expression complementation in the hybrids across genotypes and tissues for these genes, could contribute to the superior adaptation of hybrid plants to environmental stresses. The high genotype- and tissue-specific nature of non-PAV SPE and PAV SPE genes could also contribute to the diversified manifestation of heterosis across traits, environments, and genetic combinations (Flint-Garcia et al., 2009; Schnable and Springer, 2013; Marcon et al., 2017; Li et al., 2018).

These data collectively provide evidence for the different roles of non-PAV SPE and PAV SPE genes in hybrid expression complementation, and expression of the silent non-PAV SPE allele in hybrids. Results of this study shed light on the potential role of non-PAV SPE and PAV SPE genes in the manifestation of heterosis from the perspective of the transcriptome.

## Methods

### Plant Material

The inbred lines used in this study were selected to represent relevant heterotic groups in maize including the stiff stalk synthetic group (B73, B84, DKFAPW, LH145, NKH8431, PHB47, PHJ40), the non-stiff stalk synthetic group (AS5707, CR1Ht, DKHBA1, H99, LH123Ht, LH156, LH85, LH93, Mo17, NC230, NKS8326, Ny821, Oh43, PHG47, PHG39, PHW65, W64A), the iodent group (PH207, PHG29, PHG35, PHG50, PHG72, PHN11), as well as additional diverse inbred lines (Mo44, Mo45, and MoG). Hybrids were generated by crossing each of these selected inbred lines by three male genotypes that included B73 (stiff stalk synthetic), Mo17 (non-stiff stalk synthetic), and PH207 (iodent). Mo17 was also crossed with two other male genotypes, Oh43 and PHG29 (Supplemental Table 2).

Five tissues were sampled from the inbred and hybrid genotypes including seedling root at Vegetative 1 (V1), seedling shoot at V1, the middle of the eighth leaf at Vegetative 7/8 (V7/V8), the upper most elongated internode at V7/V8, and endosperm at 15 days after pollination (DAP) (Supplemental Figure 1). Seeds were planted at the Minnesota Agricultural Experiment Station located in Saint Paul, MN on 05/16/14 with 30 inch row spacing at ∼52,000 plants per hectare. Leaf and internode samples at the V7/V8 growth stage were harvested on 06/30/14 for the hybrid genotypes and 07/01/14 for the inbred genotypes. Endosperm samples were collected between 08/07/14 and 08/30/14. For the V1 tissues (root and shoot), seeds were planted on 09/03/14 into Metro-Mix300 (Sun Gro Horticulture) with no additional fertilizer and grown under greenhouse conditions (27C/24C day/night and 16 h light/8 h dark) at the University of Minnesota Plant Growth Facilities. Hybrid samples were harvested on 09/11/14 and inbred samples were harvested on 09/12/14.

Total RNA was extracted using the miRNeasy Mini Kit (Qiagen). Extracted RNA was DNase treated using the TURBO DNA-free kit (Life Technologies). Sequence libraries were prepared by the Joint Genome Institute following Illumina’s TruSeq Stranded mRNA HT preparation protocol (Illumina, San Diego, CA). Samples were sequenced on an Illumina HiSeq 2500 (Illumina, San Diego, CA) at the Joint Genome Institute to generate 150bp paired-end reads. For each RNA-Seq library 21-52 million reads were sequenced (Supplemental Table 1). Raw sequence reads have been deposited in the NCBI Sequence Read Archive (Supplemental Table 6).

### PAV Gene Identification and Filtering

PAV genes from both B73 and PH207 were obtained from Brohammer et. al (Brohammer et al., 2018) with modifications. The inbred lines in this study, including B73 and PH207, were resequenced at a depth of 12x to 65x. All genes with coverage over less than 80% of the gene model length from the B73 and PH207 resequencing data in its cognate genome B73v4 (Jiao et al., 2017) and PH207v1 (Hirsch et al., 2016) were considered recalcitrant and discarded from downstream analyses. For the remaining genes, if coverage was observed for less than 20% of the gene model the gene was classified as putatively absent in that genotype. In theory, PAV genes should not express any mRNA in any tissue in the genotype in which it is absent. However, this was observed in some instances. Thus, the PAV gene list was further filtered using the RNA sequencing results, where a PAV gene with CPM >= 1 in any tissue in a genotype in which it was classified as absent was removed from downstream analysis.

All RNA-Seq reads were mapped to only the B73v4 reference genome assembly. PAV genes in PH207 were matched to their cognate gene in B73 using a previously published gene key (Brohammer et al., 2018). As not all genes have a cognate gene pairing between assemblies, there was substantial attrition of PH207 genes in this step.

### RNA-Seq Data Processing

Reads were trimmed using Trimmomatic (Bolger et al., 2014) and mapped to the B73v4 genome assembly (Jiao et al., 2017) using the alignment software STAR with default parameters (Dobin et al., 2013). Uniquely mapped reads were assigned to and counted for the 46,117 (39,324 of them are protein-coding genes) B73v4 gene models using FeatureCounts (Liao et al., 2014). Raw read counts were then normalized by library size using the TMM (trimmed mean of M values) normalization approach to give CPMs (Counts Per Million reads) for each gene model (Robinson and Oshlack, 2010) (Available at https://de.cyverse.org/dl/d/2C79E2FB-16C5-46EA-B16F-571220169C32/Supplemental_Table7.csv). Clustering was done using the log2-transformed CPM value for each gene in each sample and “prcomp” function in R without centering and scaling (https://www.rdocumentation.org/packages/stats/versions/3.5.1/topics/prcomp). Only protein-coding genes that were expressed in at least one sample were used. Based on clustering it was determined that LH93 root tissue was incorrect and was removed from downstream analyses. For samples with multiple technical replicates, the replicate with the highest sequencing depth and mapping rate for the corresponding samples were used in downstream analyses, resulting in a total of 430 samples (Supplemental Table 1).

A gene was declared as expressed if the CPM was >= 1. An SPE gene was defined as a gene with CPM_female >= 1 & CPM_male < 0.1, or vice versa. Genes that showed expression complementation in the hybrid were defined as having CPM_hybrid > 1 or the mid-parent expression value of the parental samples. As such, the expression level in the hybrid could be as low as 0.5 and still be considered expressed to allow for additive expression behaviour. Genes that had an expression value between 0.5 and 1 in the hybrid account for only 4.2% and 9.4% of the total non-PAV SPE and PAV SPE genes. Finally, silent allele expression of non-PAV SPE genes in the hybrid was defined by having at least 10 allele specific reads.

### Non-syntenic and Curated Gene Set

The curated-gene list was downloaded from MaizeGDB with B73v4 gene IDs (https://www.maizegdb.org/gene_center/gene#gm_downloads) (Andorf et al., 2016). Synteny classifications (i.e., syntenic and non-syntenic) and assignment to maize sub-genomes were obtained from previous studies based on pairwise whole-genome alignment between maize and sorghum (Schnable et al., 2011b; Brohammer et al., 2018).

### Characterization of Allele Expression Patterns in the Hybrids

SNP genotype calls for 29 of the genotypes with resequencing data were obtained from (Mazaheri et al., 2019). For the five inbreds (DKFAPW, H99, NKS8326, W64A, Ny821) without re-sequencing data, SNPs were called by pooling all RNA-Seq reads from the corresponding inbred and mapping to the reference genome as previously described for the resequencing data (Mazaheri et al., 2019). After quality filtering, a total of 4.8 to 6.4 million SNPs were identified in each of the 29 re-sequenced inbreds relative to the B73v4 reference genome assembly, and 79-104 thousand SNPs for the other five inbreds were used to characterize allele expression patterns in the hybrids.

All hybrid reads were mapped to the B73v4 reference genome. In each hybrid sample, allelic expression differences were determined by counting RNA-Seq reads carrying at least two distinguishing SNP(s) between the two inbred parents as previously described (Zhou et al., 2019). In this method, genes with at least 20 allele specific reads and less than 10% conflicting reads were used to determine the ratio of allelic expression in the hybrid. Conflicting reads can come from reads spanning more than one variant but have contradictory predictions (i.e., a read supporting the B73 allele in one variant position and the Mo17 allele in the other variant position). Raw read counts, allele-specific read counts, source codes and scripts are available under Github repository (https://github.com/orionzhou/biomap/blob/master/Rmd/rnaseq.md).

### Genetic Distance Tree

Haplotype information for each inbred was constructed from the SNP dataset described above. A maximum-likelihood approach was used to build a genetic distance tree using IQ-TREE (Nguyen et al., 2014). Bootstrap replicates were set as 1,000 for searching the parameters. The FigTree software was used to visualize the tree (http://tree.bio.ed.ac.uk/software/figtree/).

## Author Contributions

C.N.H., conceived and designed the project. Z.L., P.Z., R.D.C. and A.B.B. performed data analysis. Z.L. and C.N.H. wrote the paper. P.Z., T.F.Z., N.D.L., C.D.H., C.R.B., S.M.K. and N.M.S. edited the manuscript. B.V., A.L. and K.B. conducted the RNA-sequencing experiment.

## Acknowledgments

We are grateful to Katie Heslip for performing RNA extractions of samples included in this study.

## Supplementary Figures

**Supplemental Figure 1.**
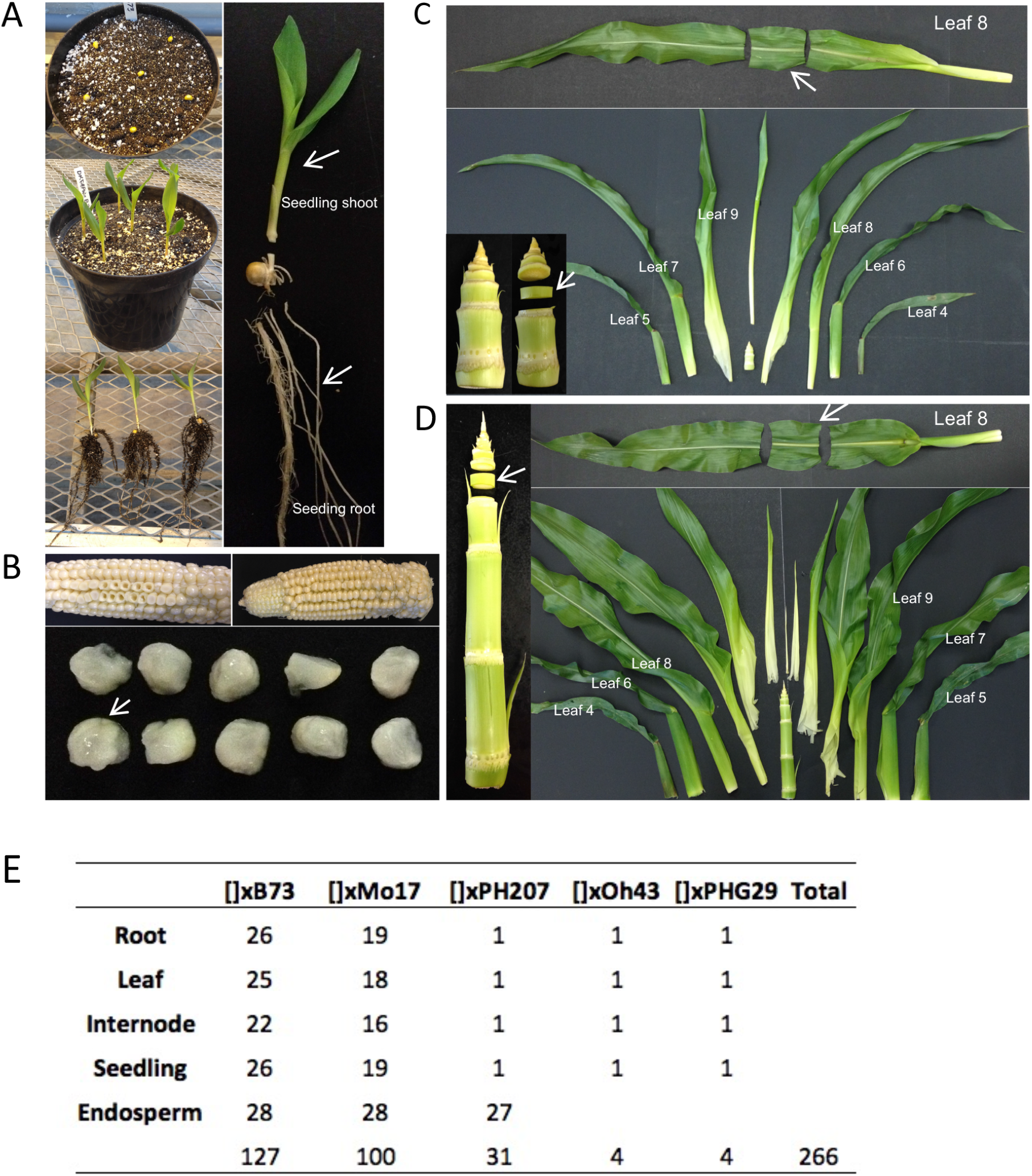
Tissues collected in this study. (A) Seedling root and seedling shoot at the V1 growth stage. (B) Endosperm at 15 days after pollination (DAP). (C, D) Leaf and internode samples at the V7/V8 growth stage for inbreds (C, V7) and hybrid (D, V8) genotypes. (E) The number of parent-hybrid triplets in each tissue across the five common male inbred parents.

**Supplemental Figure 2.**
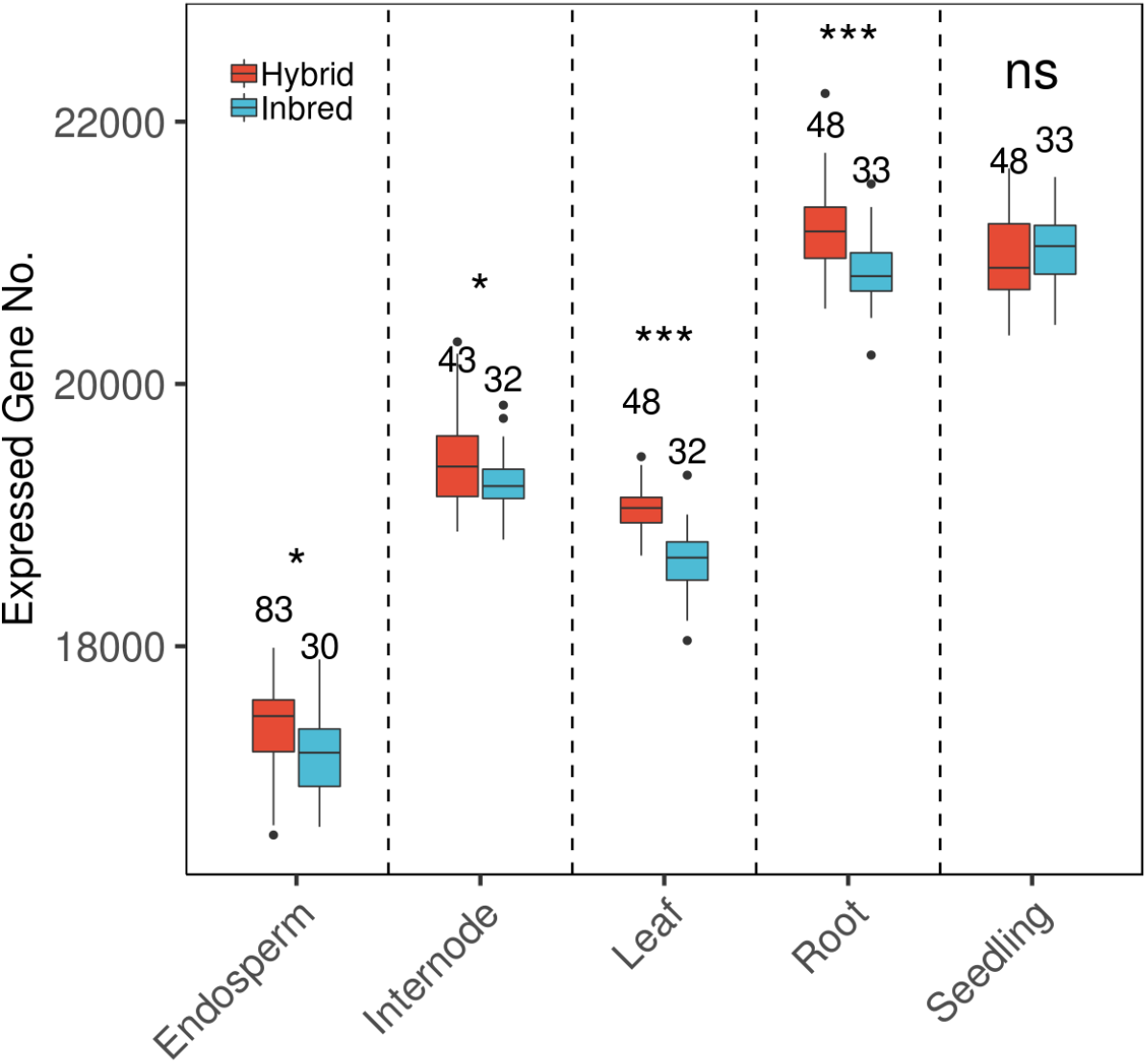
Summary of the gene expression landscape in diverse inbreds and hybrids across five tissues. Comparison of expressed gene numbers for hybrids and inbreds across tissues. The number of samples plotted in each box plot is shown above the box. Two-tail t-test for the number of expressed genes of inbreds and hybrids in each tissue was conducted to determine significant differences (“*”, p< 0.05, “***”, p< 0.0001, “ns”, not significant).

**Supplemental Figure 3.**
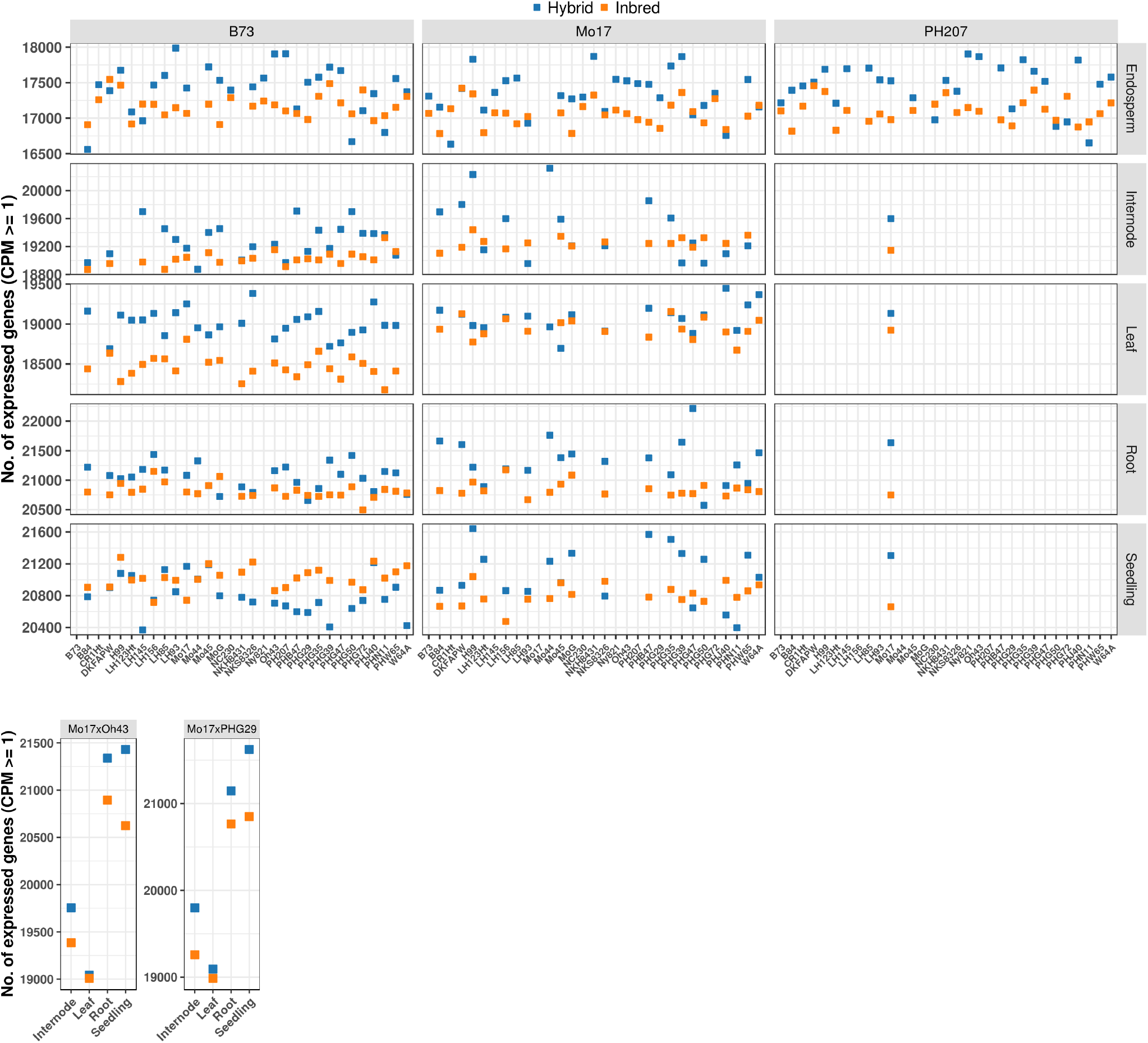
Comparison of expressed gene numbers within each parent-hybrid triplet across tissues. Comparison of expressed gene numbers for 127 Inbred x B73, 100 Inbred x Mo17, 31 Inbred x PH207 and 4 Mo17 x Oh43, 4 Mo17 x PHG29 triplets, respectively.

**Supplemental Figure 4.**
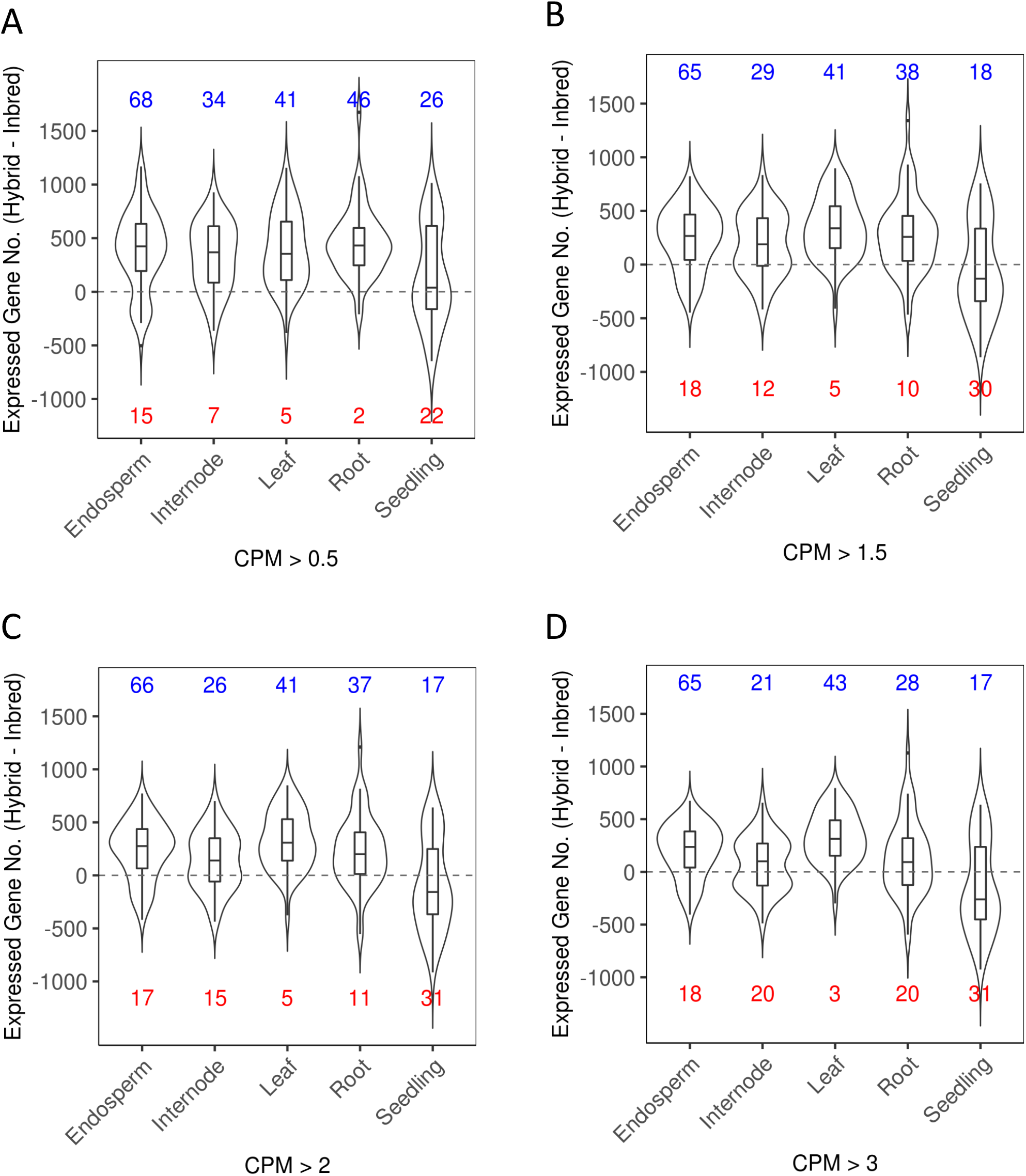
Difference in the number of expressed gene for each parent-hybrid triplet in each tissue calculated as Hybrid - 0.5*(Male-parent + Female-parent. Red numbers: the number of parent-hybrid triplets in which the expressed genes in hybrid is smaller than parent mean; blue number: the number of parent-hybrid triplets in which the expressed genes in hybrid is greater than parent mean. A-D illustrate the results when applying different criteria to declare the expression state of a gene (counts per million (CPM) > 0.5, 1.5, 2.0 and 3.0, respectively).

**Supplemental Figure 5.**
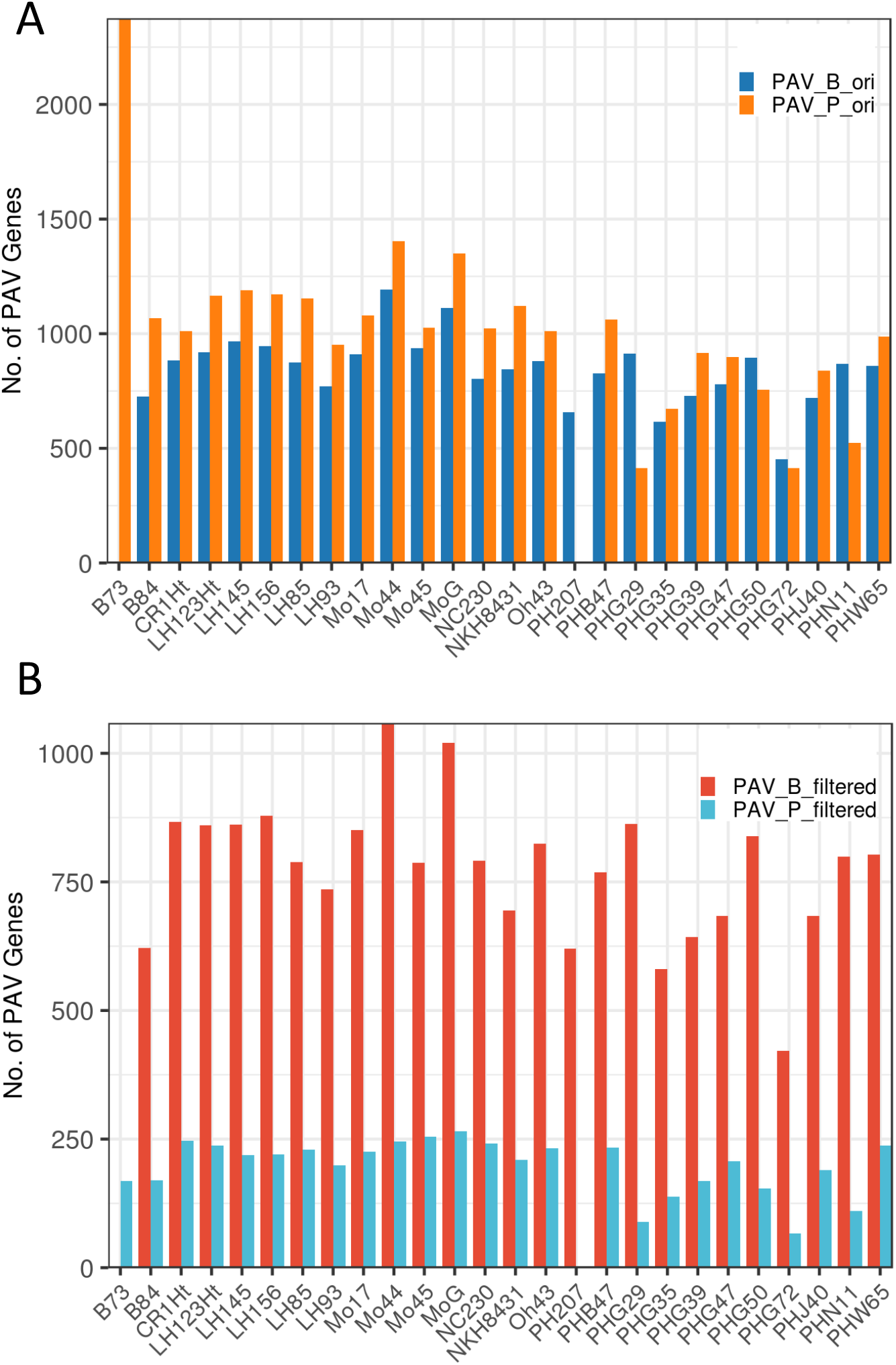
Summary of PAV genes based on alignment to the B73 and PH207 genome assemblies, respectively. (A) The number of original PAV_B and PAV_P genes identified by re-sequencing coverage for each inbred. (B) The number of PAV_B and PAV_P genes retained after the filtering using RNA-seq data (PAV genes showing evidence of expression in any tissue of the corresponding inbred were considered as non-PAV and removed from downstream analyses).

**Supplemental Figure 6.**
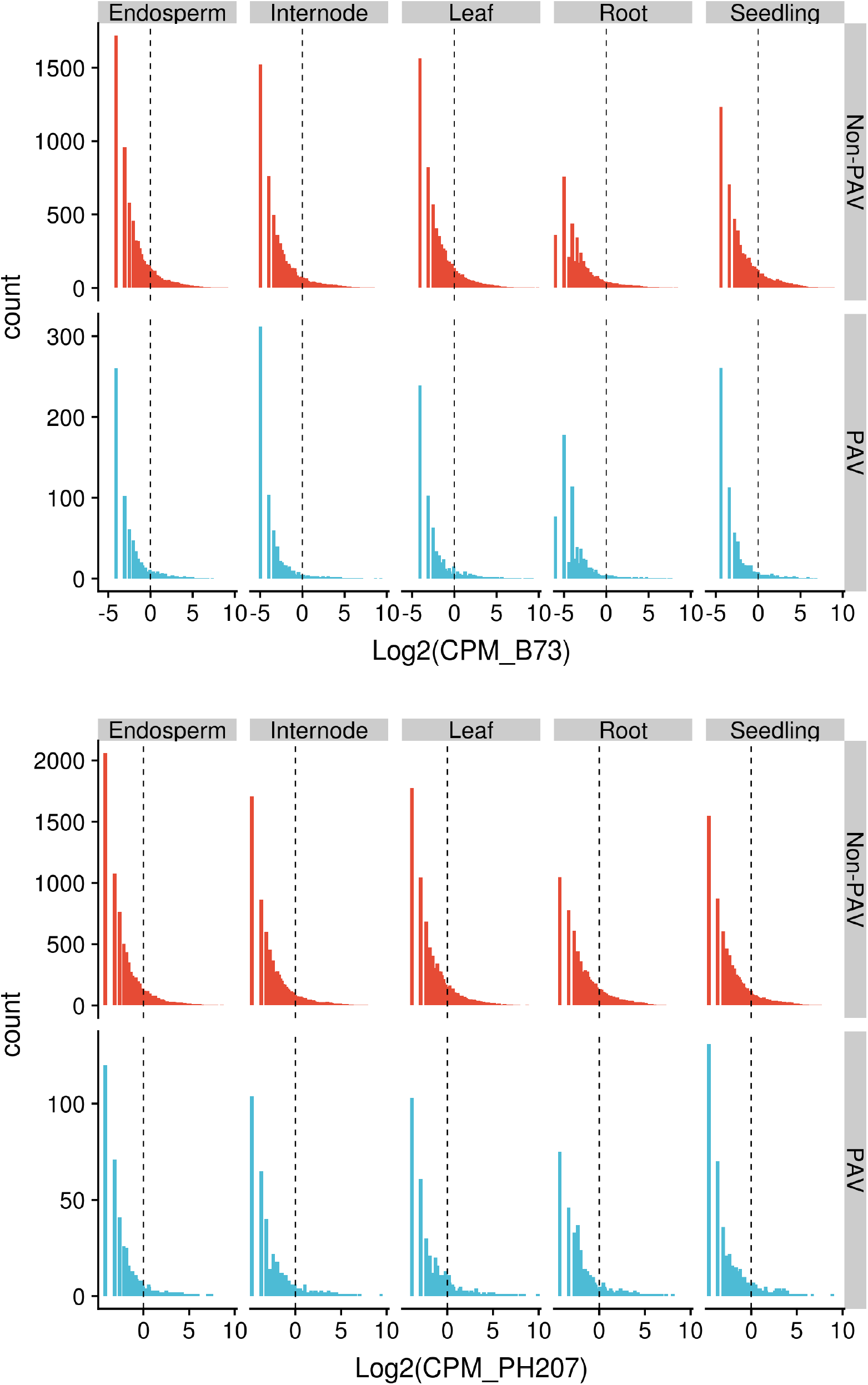
Log2-transformed expression level (CPM) of PAV and non-PAV genes in B73 and PH207. The vertical dashed line indicates the cutoff (CPM >=1) used to declare the expression status of a gene.

**Supplemental Figure 7.**
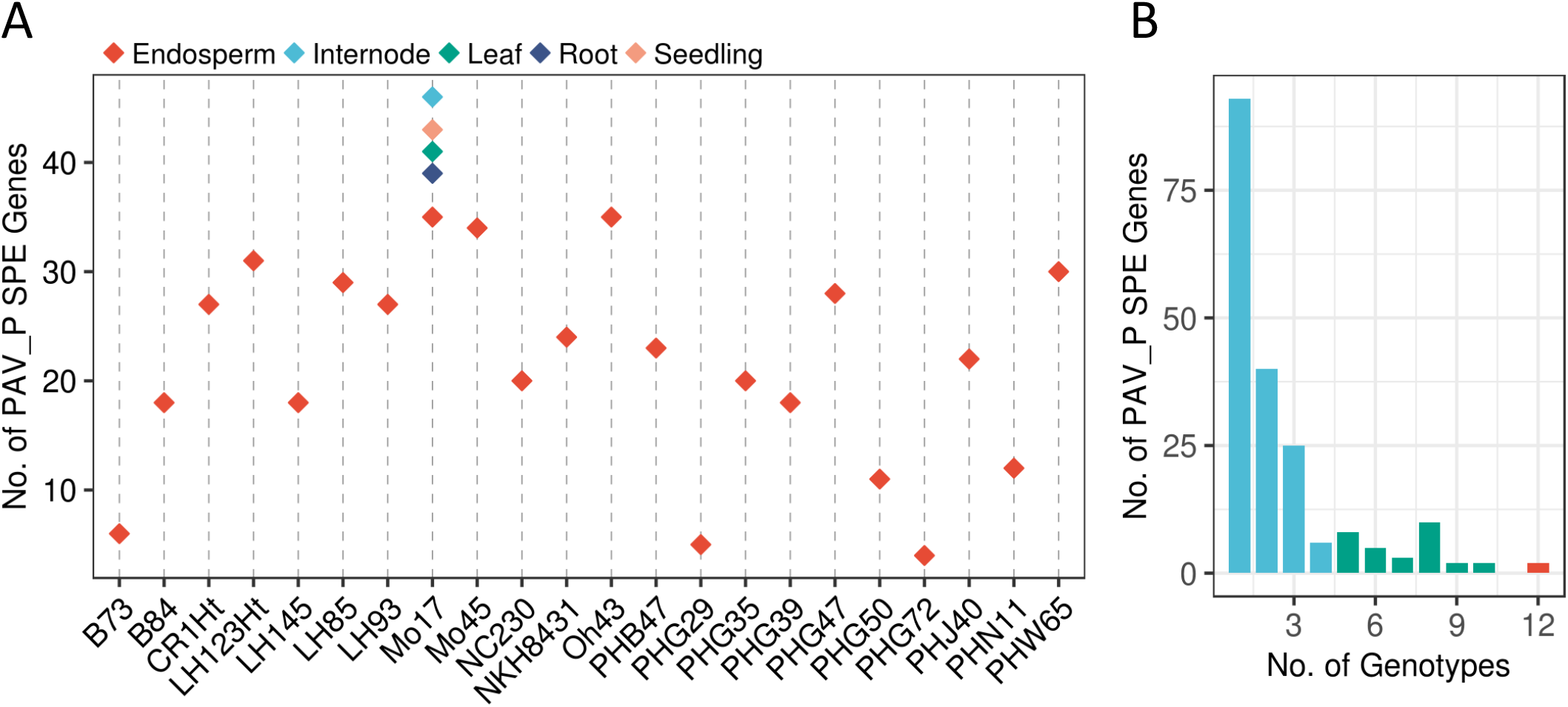
Summary of PAV SPE genes based on PH207 (PAV_P SPE). (A) The number of PAV_P SPE genes for each female parent and tissue. (B) The number of PAV_P SPE genes detected in 1-22 female parents. Each bar represents the number of PAV_P SPE genes shared by how many inbred parents (x-axis). Light blue, green, and red color of bars represent the number of PAV_P SPE genes shared across <20%, 20%-50% and >50% of the total inbred lines respectively.

**Supplemental Figure 8.**
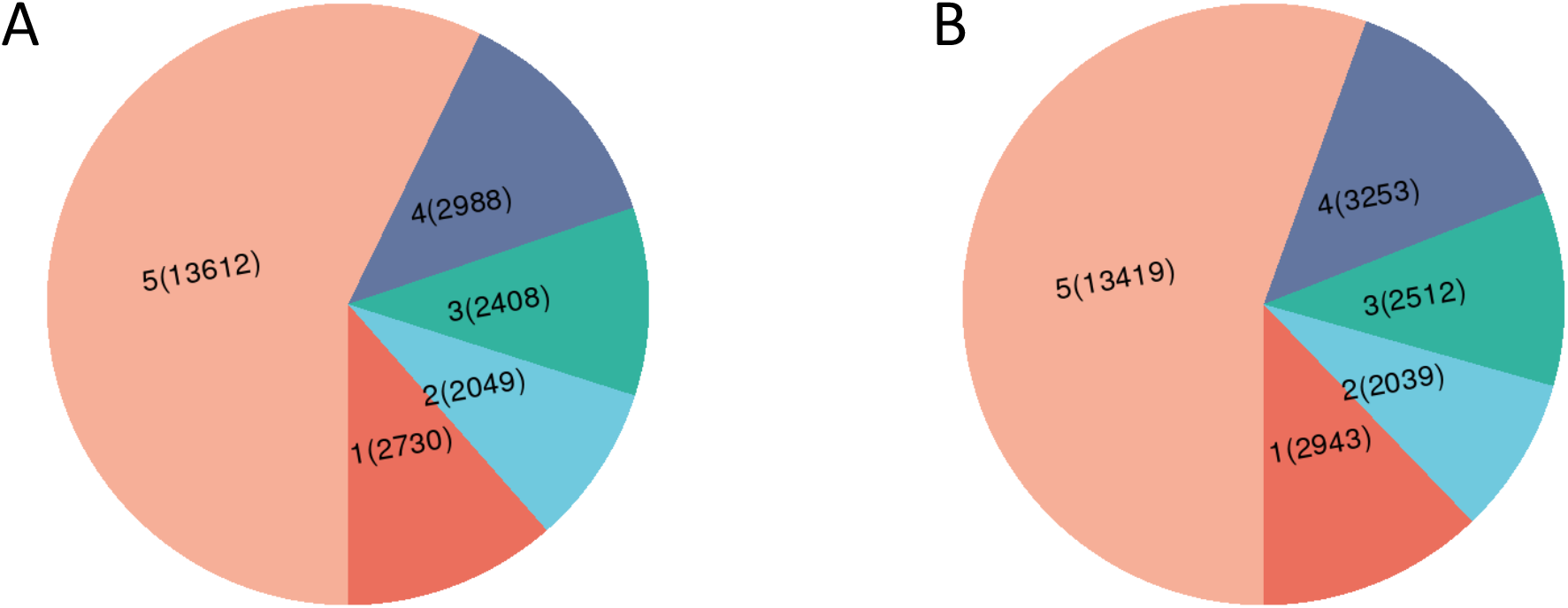
Tissue-specificity of non-PAV genes in B73 (A) and PH207 (B). Genes expressed in at least one of the five tissues in B73 or PH207 were used to conduct this analysis. Each part of the pie chart represents the number of non-PAV genes (number in parentheses) shared by the number of tissues (1-5, outside parentheses).

**Supplemental Figure 9.**
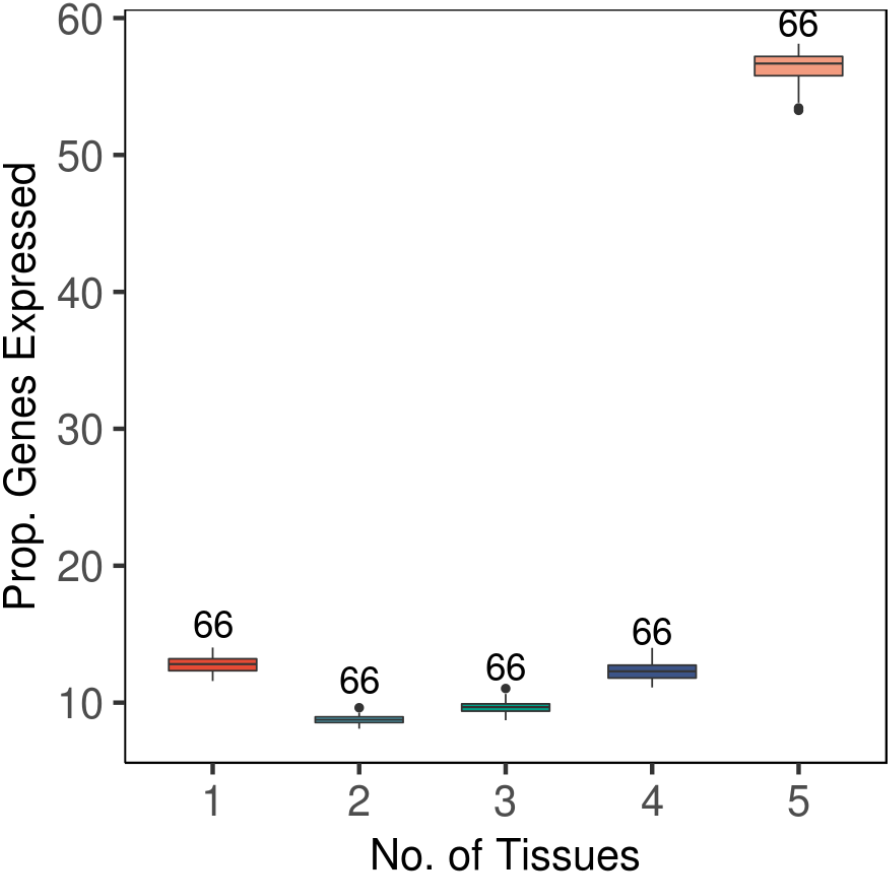
Tissue-specificity for genotypes that were sampled for all five tissues. Numbers above each box are the number of genotypes. Over half of the expressed genes are expressed in all the five tissues.

**Supplemental Figure 10.**
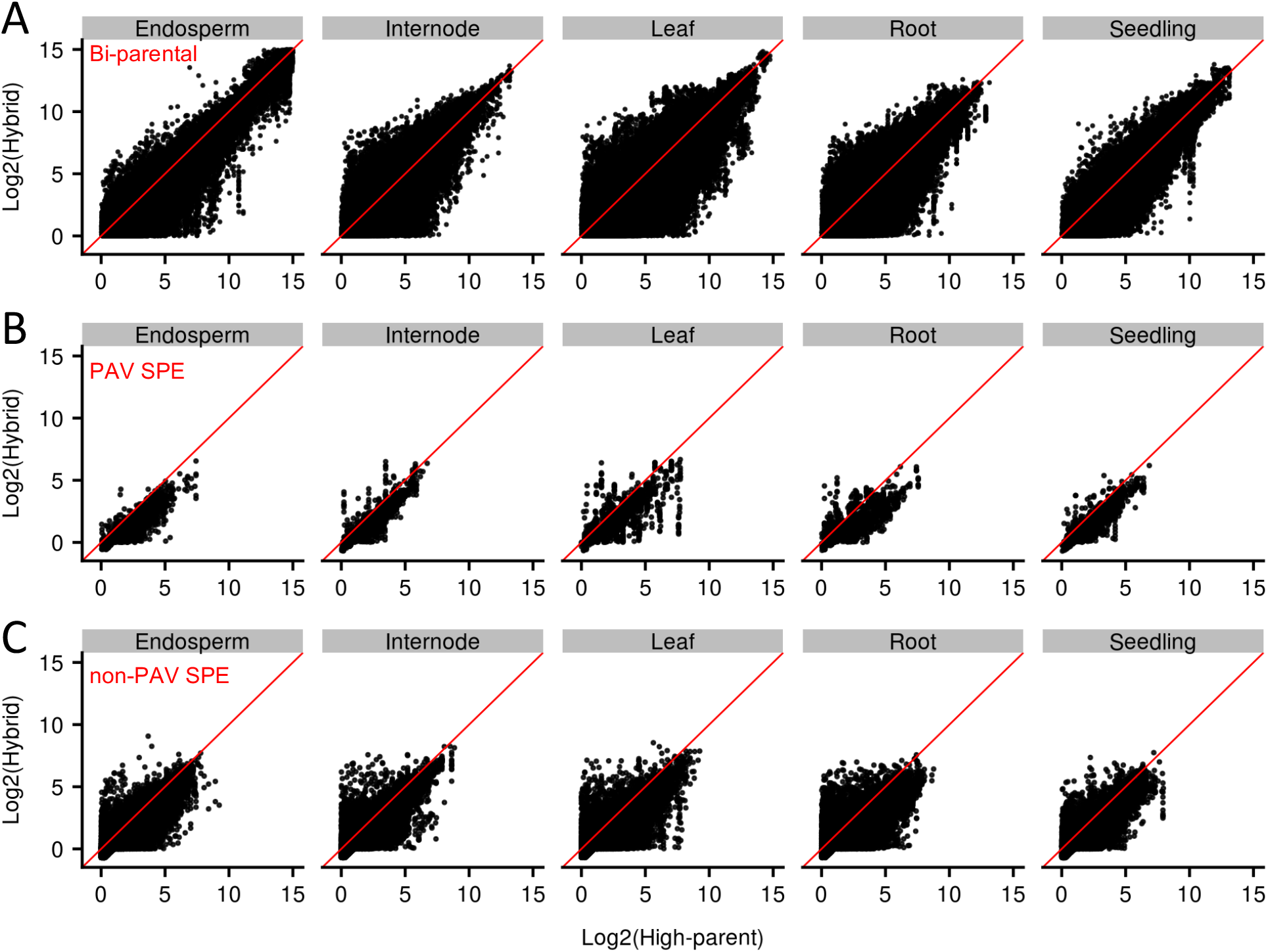
Expression level of genes in different expression pattern categories. The log2-transformed expression level (CPM) of each gene in the high-parent inbred and the hybrid was plotted for each pattern. Dots that fall on the red line indicate genes that have the same expression level in the hybrid and the high-parent.

**Supplemental Figure 11.**
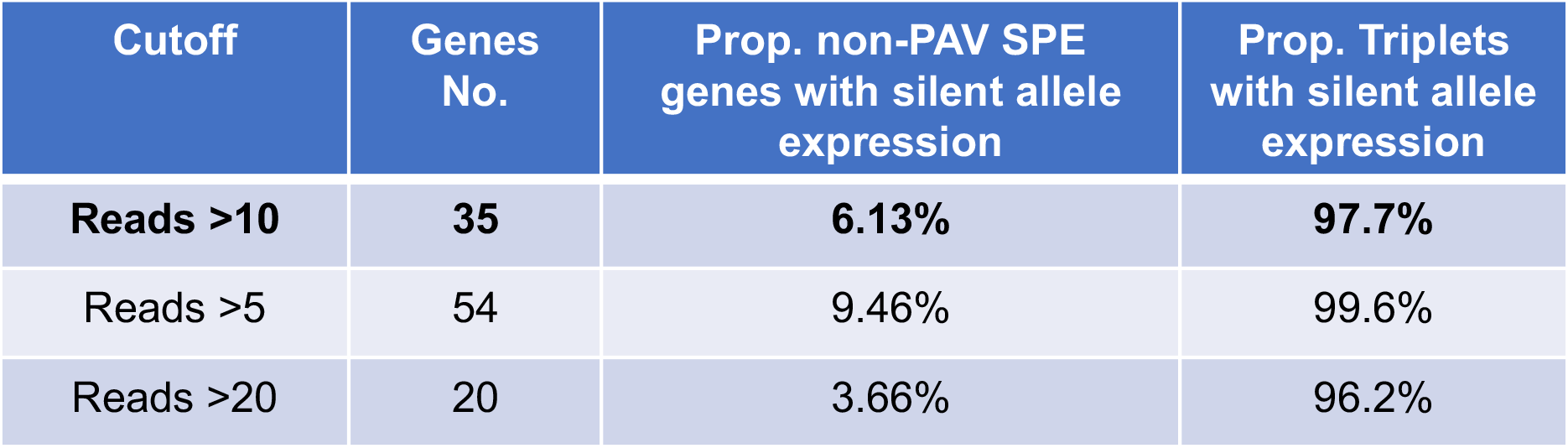
Evaluation of silent allele expression in the hybrid determined using different cutoffs.

**Supplemental Figure 12.**
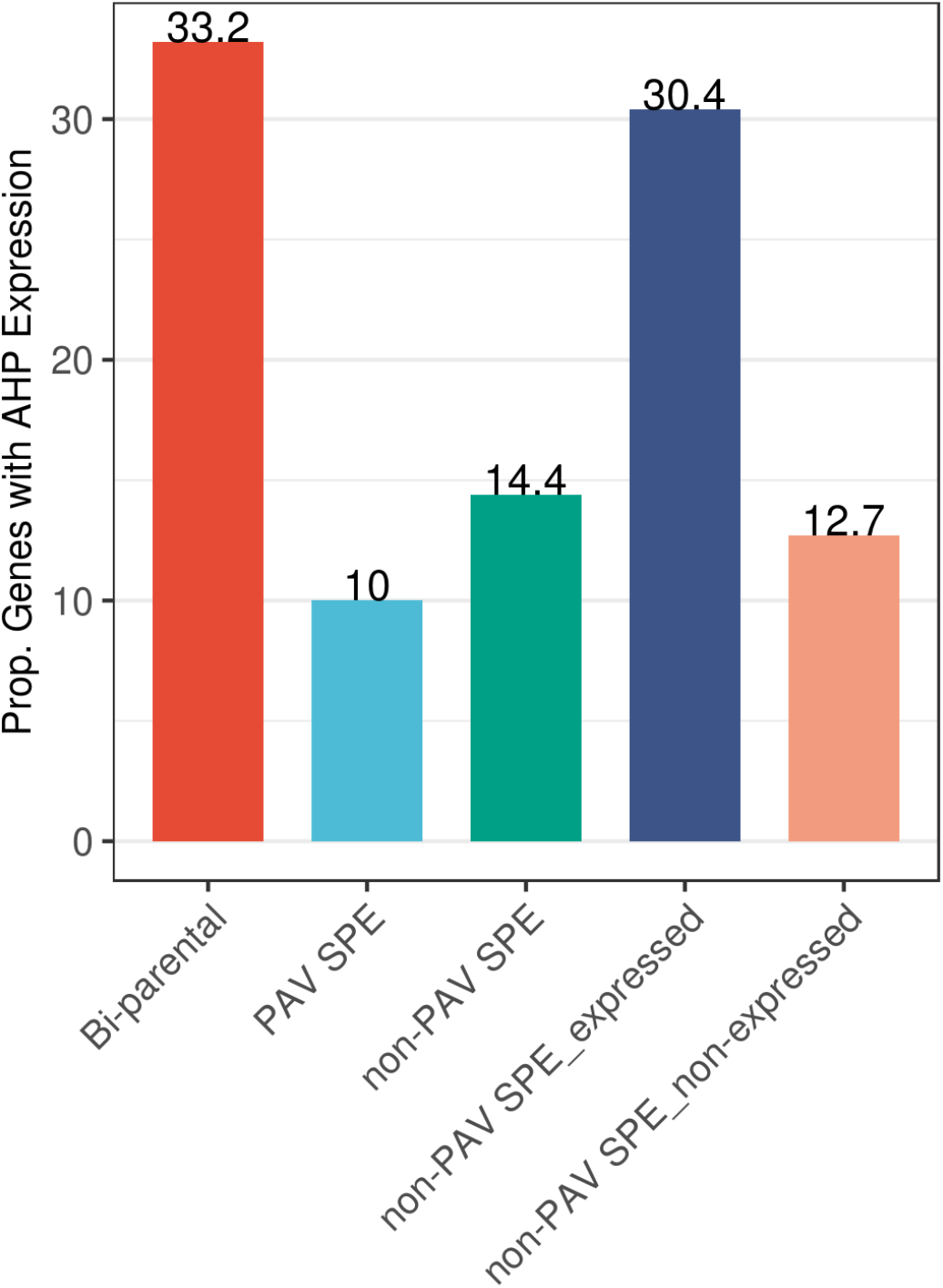
Proportion of genes in the hybrid that showed above-high parent (AHP) expression level for each pattern (averaged across samples). The dark blue and orange bar indicate non-PAV SPE genes with and without silent allele expression in the hybrid, respectively.

**Supplemental Figure 13.**
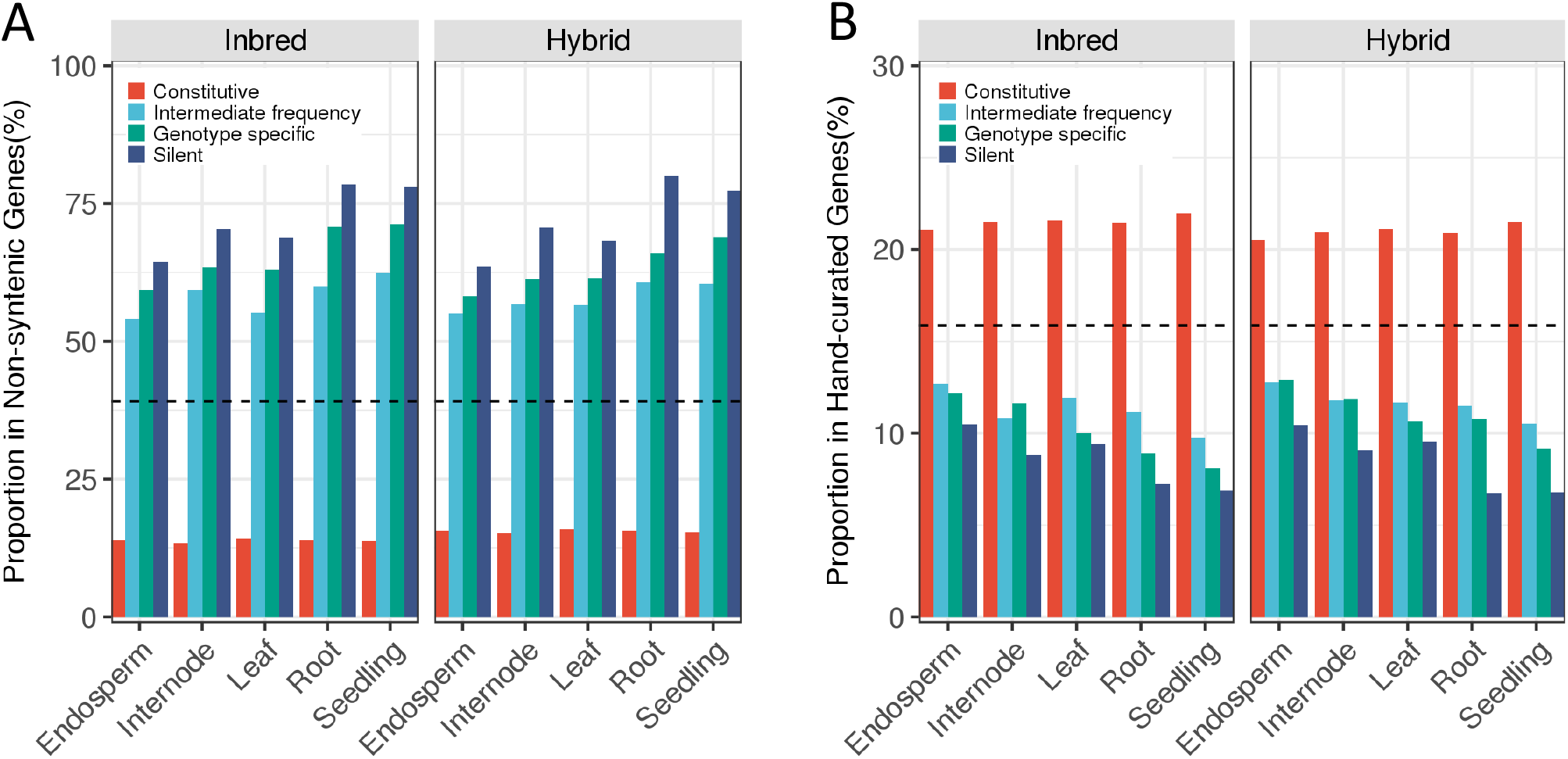
Proportion of genes in each expression category defined in Figure 2A and 2B that are in the non-syntenic and curated gene sets. (A) The proportion of genes in each expression category that are non-syntenic across tissues in inbreds and hybrids, respectively. Black-dashed line shows the genome-wide proportion of non-syntenic genes in the maize genome (39.2%, 15,402/39,324). (B) The proportion in curated gene list genes in each expression category across tissues in inbreds and hybrids, respectively. Black-dashed line shows the genome-wide proportion of curated gene in the maize genome (15.9%, 6,239/39,324).

